# A DNA-barcoded Luria-Delbrück assay resolves mechanisms of adaptation

**DOI:** 10.64898/2026.08.03.742544

**Authors:** Kabir Husain, LaNijah Flagg, David Pincus, Arvind Murugan

## Abstract

When populations encounter a harsh new environment, they adapt in many different ways – from spontaneous genetic changes to transient non-genetic plasticity. Distinguishing between mechanisms of adaptation is often difficult, as the key events are rare and hard to observe directly. Here, we introduce BarFly, a Luria-Delbrück assay that uses DNA barcoding and deep sequencing to resolve the Luria-Delbrück distribution – the statistics of adaptation to a new environment – over four orders of magnitude. At this statistical depth, the shape of the distribution encodes the underlying adaptive mechanism. We find that DNA mismatch repair mutants shift the distribution’s scale without changing its shape, indicating their only effect is as mutator alleles. A pulse of UV mutagenesis, however, adds a second, statistically distinguishable mode atop the spontaneous background, showing that different mechanisms of adaptation can be quantified directly from the distribution. By sampling rare-event statistics with DNA barcodes, BarFly provides a general method for studying mechanisms of adaptation in evolving populations from microbes to cancers.

## I. INTRODUCTION

To adapt to unfamiliar environments, evolving populations rely on a supply of heritable variation. The form that heritable variation takes depends on the context, ranging from spontaneous changes to genetic material – e.g., *de novo* mutations, recombination, horizontal gene transfer [1] – to stress- or drug-induced mutagenesis [2], or even non-genetic variation in the form of sustained fluctuations in gene expression [3–7] or the inheritance of protein prions [8, 9]. Determining how a given population generates heritable variation is often a key aspect of predicting its future trajectory in the lab, or intervening in that trajectory for pathogens and cancers in the clinic. Yet this is difficult to measure directly, as the causative variants arise as rare stochastic events at low frequencies in the population.

Eighty years ago, Luria and Delbrück [10] proposed an alternative, statistical approach to study the nature of heritable variation. By quantifying the number of mutants arising in replicate bacterial cultures, they inferred that mutations arise spontaneously and even measured the mutation rate – a full decade before the structure of DNA was determined. Since then, their ‘fluctuation assay’ has been celebrated as a landmark experiment [11], used to illustrate the power of quantitative thinking in biology [12, 13], and routinely employed to measure rates of mutation [14–17] and other rare, heritable processes [18, 19]. More recent work has used the assay to study drug resistance in cancers and fungi [3, 7], the transient heritability of gene expression [4], genetic variation in biofilms [20], driver mutations in cancers [21], and the mutation rate of persister cells after drug treatment [22].

However, the modern applicability of the Luria-Delbrück assay is fundamentally limited by the number of statistical replicates used in the experiment (Fig. 1a). Most studies use 50-100 replicates, with Luria and Delbrück themselves doing 87 [10]: enough to support the existence of spontaneous mutations, but, as later analysis showed, insufficient statistical power to rule out CRISPR-like induction mechanisms [23]. By contrast, over fifty years of theory [24] argue that many phenomena – such as partially heritable phenotypic states [25– 27], phenotypic delays following mutation [28, 29], or heterogeneous mutation rates within a clonal population – could be identified if the Luria-Delbrück distribution were sufficiently well resolved. Recent efforts using robotics reach *∼* 10^2^ − 10^3^ replicates [14, 31], while theory suggest that *>* 10^4^ replicates may be needed to reliably sample the rare events that distinguish biologically distinct hypotheses [23].

**FIG. 1.**
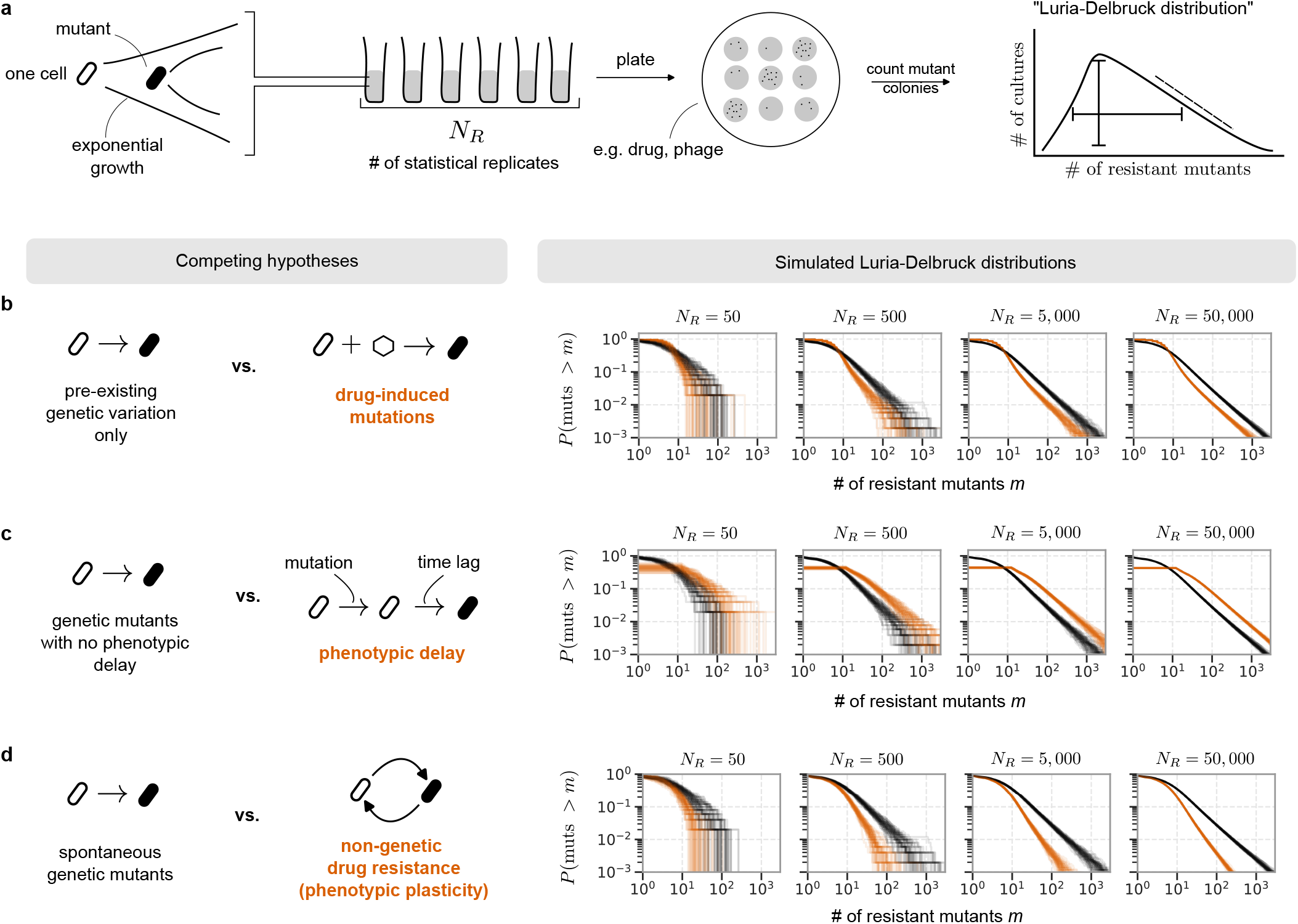
Distinguishing between adaptive mechanisms requires deep sampling of rare events in the Luria-Delbrück distribution. **(a)** Schematic of the Luria-Delbrück experiment. The key experimental parameter is the number of replicate cultures tested, *N*_*R*_, which determines the statistical resolution of the distribution. **(b, c, d)** Overlaid Luria-Delbrück distributions from 50 simulated experiments with the indicated number of statistical replicates *N*_*R*_. Black curves show the canonical Luria-Delbrück distribution (i.e. adaptation via spontaneous genetic mutations only), while orange curves include (b) drug-induced resistance, (c) a phenotypic delay after mutation, or (d) phenotypic adaptation via transient, reversible cell states.

Here, we use DNA barcodes to sample from the Luria-Delbrück distribution at a high statistical resolution. Termed BarFly, by this approach we achieve *>* 10^5^ replicates in a one-pot experiment, resolving the Luria-Delbrück distribution over several orders of magnitude. We show experimentally – using mismatch-repair mutations and UV-induced mutagenesis – that different mechanisms of adaptation can be resolved as a consequence: allowing their relative quantification, and classifying genetic and environmental perturbations by their impact on different parameters of evolution.

## II. RESULTS

### A. The shape of the Luria-Delbrück distribution encodes mechanisms of adaptation

The Luria-Delbrück experiment quantifies the number of mutants in a population grown from a single wildtype cell. Typically, this is obtained by plating the entire population onto solid media with a selective agent (e.g. a drug or phage) that kills the wildtype. The few mutants that survive appear as colonies. The process is stochastic, and so the experiment is performed in replicate to sample from the so-called Luria-Delbrück distribution, Fig. 1a.

Luria and Delbrück’s insight was that the quantitative form of the distribution depends on the mechanisms by which mutants arise. For instance, if mutations occur spontaneously, then they occasionally occur early in population growth and give rise to a large mutant lineage. In the simplest, canonical model, as analysed by Lea and Coulson [32], Haldane [33], Mandelbrot [34], and others [35, 36], these ‘jackpots’ endow the Luria-Delbrück distribution with a long-tailed power-law decay, *P* (*m*) *∼ m*^−2^ – which, in particular, results in a large variance used by Luria and Delbrück to reject the hypothesis of induced mutagenesis [10]. In more complex scenarios, many years of theoretical work [24] has shown that multiple additional processes shape the quantitative form of the distribution. For instance, mutants that decrease replicate fitness produce smaller jackpots, lead to a steeper power-law *P* (*m*) *∼ m*^*−α*^, *α >* 2 [36]. In some contexts, mutations can occur both spontaneously as well as induced by stress or (in phage resistance) by CRISPR-like immunity, leading to a complex distribution *P* (*m*) that convolves the Luria-Delbrück power-law with a Poisson distribution [14, 23].

These considerations naturally lead to the inverse problem: under what conditions can the underlying processes be inferred from an empirically observed Luria-Delbrück distribution? In Fig. 1b, c, d we present simulations of Luria-Delbrück assays with different mechanisms discussed in the literature: (b) stress- or drug-induced mutagenesis [2, 37, 38], (c) a time delay between mutation and when the mutant phenotype is expressed (e.g., due to protein expression or degradation) [28], and (d) drug resistance arising not from genetic mutations but instead transiently heritable phenotypic states [3, 7, 25]. In these simulations, we varied the number of statistical replicates *NR* used in the experiments, while using parameter values typical for biological adaptation, see SI for details.

We find that, in all cases, resolving the distributions that correspond to biologically distinct hypotheses requires *NR >* 10^3^. Note that Luria and Delbrück and most following works use *NR ≈* 50 − 100, a statistical resolution at which distributions are poorly distinguishable. The current state-of-the-art using either robots or long multi-day experiments does achieve *NR ≈* 1000 (720 in [14], 1104 in [39], 1920 in [22]) with some effort, a scale at which the power-law nature of the Luria-Delbrück distribution starts to become apparent. Only when the number of statistical replicates crosses thousands are rare events resolved with sufficient statistical power to distinguish between biologically distinct hypotheses.

### B. BarFly: Resolving the shape of the Luria-Delbrück distribution with barcode sequencing

We therefore sought to develop a high-resolution Luria-Delbrück assay capable of the statistical power necessary to resolve different adaptive mechanisms. To do so, we replaced physically-separate replicate cultures with DNA-barcoded lineages, and colony counting by deep sequencing, Fig. 2, reasoning that the number of mutants produced by each lineage could be inferred from barcode frequencies seen after selection – with each barcoded lineage acting as an independent sample from the Luria-Delbrück distribution. The main advantage of this approach is that the number of statistical replicates would be only limited by the efficiency of integrating DNA barcodes into a population, which is easily in excess of 10^5^ in many systems [40–45]. We term this approach a barcoded fluctuation assay, or *BarFly*

**FIG. 2.**
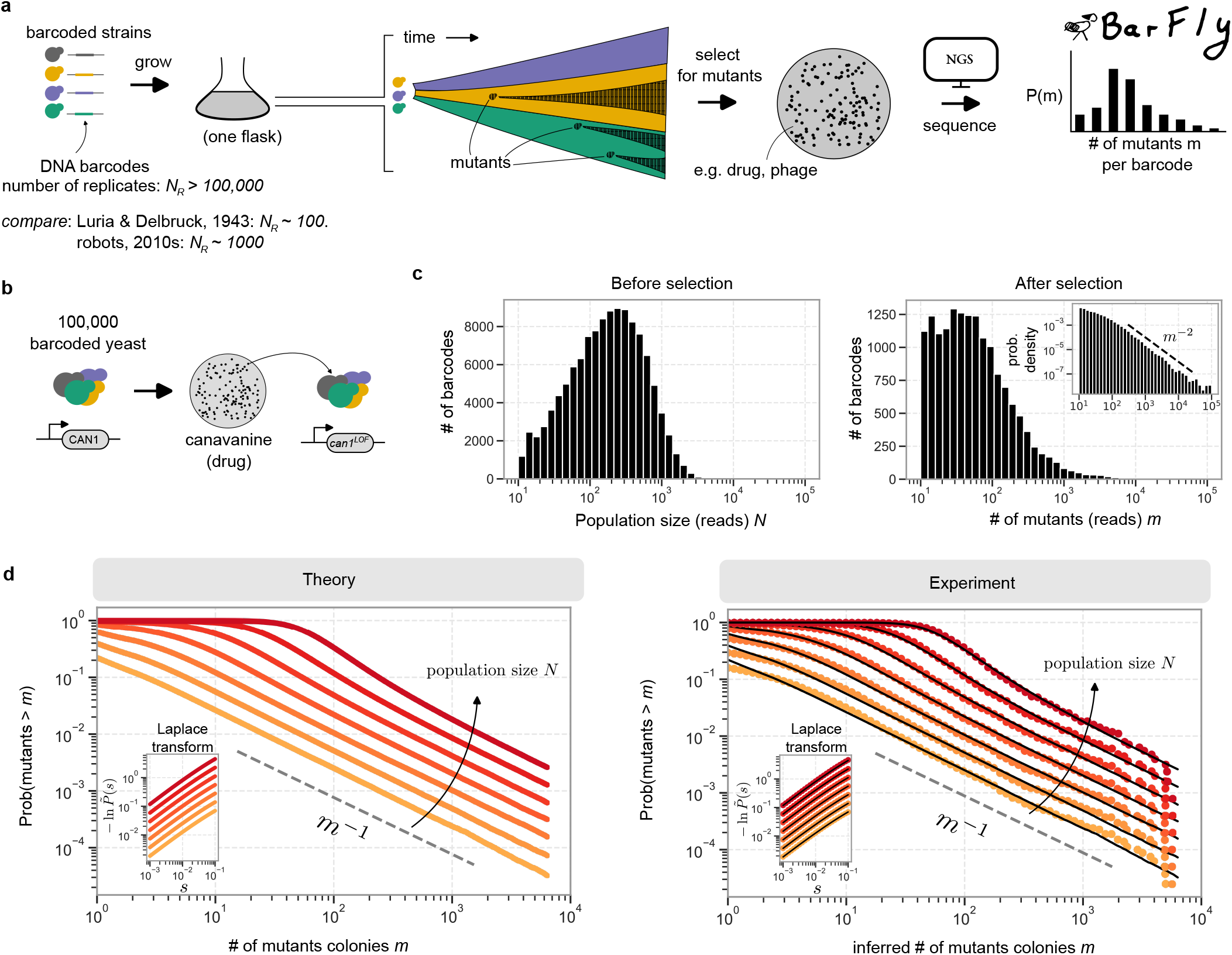
BarFly: high-resolution sampling from the Luria-Delbrück distribution with DNA barcodes. **(a)** Traditionally, Luria-Delbrück assays are performed with physically separate replicate cultures. In BarFly, replicates are instead DNA barcoded and grown together, with colony counting replaced by deep sequencing to track individual replicates. **(b)** We applied BarFly to the acquisition of canavanine resistance in budding yeast, which occurs via loss-of-function mutations in the can1 gene. **(c)** The distribution of reads per barcode before (left) and after (right) selection. Inset shows the probability density of reads per barcode on logarithmic axes; dashed line has exponent -2. **(d)** Comparing experiment and theory: plots show (Left) simulated and (Right) experimental Luria-Delbrück distributions at varying population size *N*. Lineages of varying *N* were obtained from a single experimental dataset by computationally merging barcodes together. Dashed line has exponent -1, and overlaid black lines on the right are fits to the Luria-Delbrück distribution. Insets show the Laplace transform (also known as the cumulant generating function), with fits on the right-hand-side inset in black; see text for details.

To test the resolving power of BarFly, we turned to a standard Luria-Delbrück assay: the selection of canavanine resistant mutants of *S. cerevisiae*, Fig. 2b, a selection scheme often used to measure mutation rates in yeast [14, 16]. A 12-nucleotide barcode was integrated into the trp1 locus of a lab yeast strain (BY4741 msh2Δ) at an estimated diversity of 10^5^ barcodes. A million cells (10× cells/barcode) were grown neutrally for 13 generations to *∼* 10^10^ cells, and then mutants were selected for by the addition of the drug canavanine.

We tracked barcode frequencies by deep sequencing over time. Before selection the distribution was well peaked (Fig. 2c, left; median 5 × 10^4^ cells per barcode). After selection it was markedly broad and power-law distributed (Fig. 2c, right and inset): of the 119, 672 bar-codes tracked, 84% were undetectable – indicating a failure to produce canavanine-resistant mutants – while the 100 most frequent accounted for 45% of reads. This post-selection distribution was stable for at least 16 generations (SI Fig. S1g), indicating the shift in barcode frequencies was driven by canavanine selection rather than other evolutionary processes.

We sought to compare the measured barcode-frequency distribution with the theoretical Luria-Delbrück distribution. Loss-of-function mutations in *can1*, which confer canavanine resistance, are expected to arise spontaneously and be neutral in the absence of the drug, satisfying the assumptions of the canonical Luria-Delbrück distribution. Although the distribution has no closed form, for large *m* it decays as *P* (*m*) *∼ m*^−2^ [36], consistent with our data (right inset of Fig. 2c). A very good approximation is available for its cumulant generating function [35], also known as its Laplace transform,

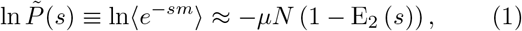

where *µ* is the mutation rate and *N* the population size prior to selection (see SI for the derivation and a definition of the generalised exponential integral *Ep*(*z*)).

To compare data with theory, we converted reads per barcode to absolute mutant numbers using a total mutant count *M* estimated by colony counting and refined during fitting, and – because our barcoded lineages vary in pre-selection size N (Fig. 2c, left), unlike the fixed-*N* assumption of the theory – computationally merged barcodes into virtual lineages spanning a range of *N*, justified by the stability of the Luria-Delbrück distribution under addition [34, 36]. Empirical distributions and their cumulant generating functions were then fit by the method of least-squares (SI).

In Fig. 2d we present the results of this analysis. Each experimental cumulative distribution, probed at different population sizes *N*, exhibits the 1*/m* power-law tail characteristic of the cumulative of the Luria-Delbrück distribution. The cumulant generating function computed from the data fits well to the analytical form of Eq. 1 (Fig. 2d right, inset). Fits of both the cumulative distribution *P* (muts *> m*) as well as its cumulant generating function consistently returned a mutation rate of *µ ∼* 2 × 10^−6^ per generation, with 95% confidence intervals (CIs), obtained by bootstrapping, of [1.5, 2.9] × 10^−6^ from fitting the cumulative, and [1.7, 2.4] × 10^−6^ from fitting the cumulant generating function, see Fig. S4f. Using a previously determined value of 236 basepairs as the mutational target size for canavanine resistance [14], we obtain a per-basepair mutation rate of *µ*bp *∼* 9 × 10^−9^ substitutions per basepair per generation, a value consistent with published values for strains with the msh2Δ genotype used here [17, 46–48].

With this single value of *µ* = 2 × 10^−6^, we compute and overlay simulated Luria-Delbrück distributions on-to the data (black lines in Fig. 2d). We observe a good fit at all *N*, and conclude that BarFly faithfully samples from the Luria-Delbrück distribution, achieving statistical resolution that spans four orders of magnitude in both the number of replicates as well as mutant colony number.

### C. Mismatch-repair mutations reshape the Luria-Delbrück distribution only through the spontaneous mutation rate

We next sought to assess how the Luria-Delbrück distribution, as resolved by BarFly, responds to genetic perturbations that alter the parameters of evolution. We chose to first characterise mutants of the DNA mismatch repair (MMR) pathway, whose loss-of-function variants have been reported to globally increase the mutation rate [46, 49], are associated with genetic instabilities in cancers [50–52], and are widely used to generate mutator strains in yeast [53–55]. We barcoded wildtype, msh2Δ, and msh6Δ strains to a diversity of *>* 10^4^ barcodes each, and subjected each to a BarFly assay as in Fig. 2: selecting on canavanine, and sequencing barcodes before and after selection.

In Fig. 3b we present the Luria-Delbrück distributions for each strain, analysed and fit as in Fig. 2. Each shows the characteristic 1*/m* power-law decay expected of the canonical Luria-Delbrück distribution. Further, all three data-sets can be fit by varying only the spontaneous mutation rate *µ* in the Luria-Delbrück distribution. The value of *µ ∼* 2.0 × 10^−6^ (95% CI: [1.8, 2.3] × 10^−6^) obtained for msh2Δ in this experiment is in-line with the (independently obtained) data of Fig. 2, as well as with prior published values. The inferred mutation rates for the wildtype (*µ ∼* 4.3 × 10^−7^, 95% CI: [3.5, 5.2] × 10^−7^) and msh6Δ (*µ ∼* 1.1 × 10^−6^, 95% CI: [0.9, 1.4] × 10^−6^) are also within the range of values previously reported [14, 47, 53, 56].

**FIG. 3.**
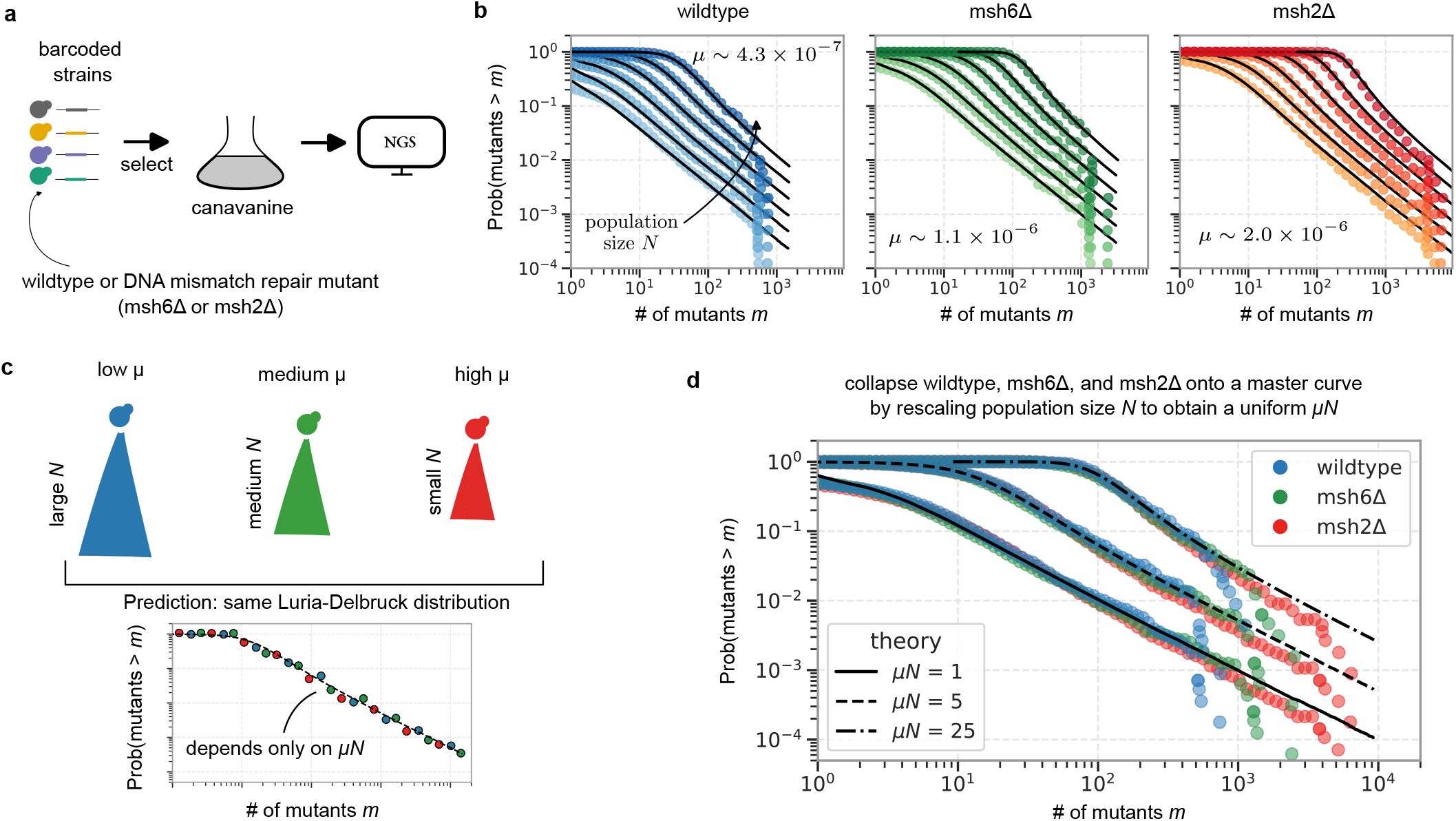
The Luria-Delbrück distributions of mismatch repair mutants can be collapsed onto a master curve by rescaling population size *N*. **a** Schematic of the experiment, in which we performed a BarFly assay on wildtype, msh6Δ, and msh2Δ strains. **b** Distributions of mutant number per lineage for each of the three strains. As before, lineages of varying *N* were obtained by computationally merging barcodes together. Black lines are computed Luria-Delbrück distributions with the indicated value of mutation rate *µ*. **c** The Luria-Delbrück distribution depends on *µ* only through the product *µN* ; if strains differ only in their spontaneous mutation rate *µ*, their distributions will be identical if population size *N* is rescaled to equalise *µN*. **d** Data collapse of the Luria-Delbrück distributions of all three strains onto each other by rescaling *N* to achieve a target *µN* = 1, 5, or 25. Black lines are simulated Luria-Delbrück distributions. Data collapse indicates that the mismatch-repair mutants affect only the availability of pre-existing mutations by modulating the spontaneous mutation rate *µ*.

If these three strains differ only in their mutation rates *µ* and not in other adaptive aspects such as those suggested in Fig. 1, then the distributions should depend only on the overall scale parameter *µN* (Eq. 1) – and should collapse onto each other when the population size *N* is appropriately rescaled, Fig. 3c. Using the fitted values of *µ* for each strain, we chose population sizes *N* for each strain to obtain *µN* = 1, 5, or 25. We then combined barcodes to form virtual lineages with the target value of *N*, and plotted the resulting Luria-Delbrück distributions in Fig. 3d.

We observe that the distributions from the three different strains collapse onto a single master curve for each value of *µN*, agreeing well with the canonical Luria-Delbrück distribution at that value of *µN* and indicating that the three strains differ only in their rate of spontaneous mutagenesis: changes to any additional adaptive mechanism (such as those outlined in Fig 1) would have deformed the shape of the LD distribution and the data would not have collapsed in Fig. 3d.

### D. BarFly quantifies the relative amounts of spontaneous and induced drug resistance

We then sought to go beyond spontaneous mutations and test whether BarFly can identify other modes of adaptation. Recent work argues that drug-resistant mutations in microbes and cancers [2, 37, 38] can be induced by stressful conditions – often the drug itself, with antimicrobials such as ciprofloxacin directly targeting the DNA replication machinery, though other, less direct molecular mechanisms exist [57]. In another context, while Luria and Delbrück famously ruled out induced mutations in the acquisition of phage resistance in bacteria, we now know that phage resistance can be induced via CRISPR-based immunity [23]. In each case, the induced mutants are added on top of the spontaneous background (Fig. 4a), distorting the Luria-Delbrück distribution away from its single-parameter form [14, 23]. We reasoned that the statistical power of BarFly could identify this distortion and quantify the relative extent of spontaneous and induced mutations.

**FIG. 4.**
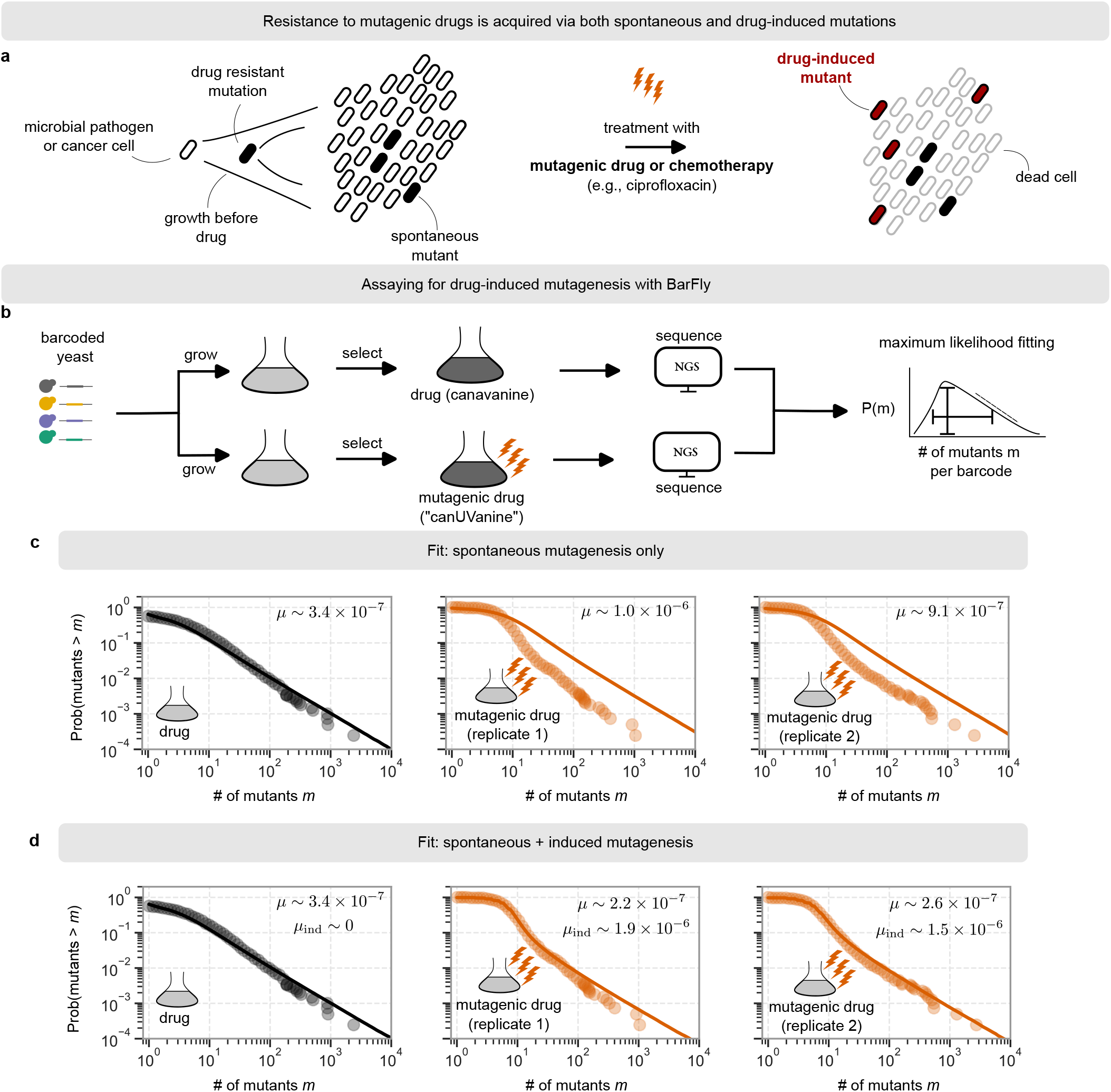
BarFly quantifies the relative contribution of spontaneous and induced mutagenesis. **a** Some drugs are reported to induce mutations in microbial pathogens and cancers. These induced mutations co-exist with spontaneous mutations that arose before treatment with the drug. **b** We assay for induced mutations by a BarFly experiment in which barcoded populations are exposed either to canavanine as before, or a compound selection we term ‘canUVanine’ that mimics drug-induced mutagenesis via a brief exposure to ultraviolet light at the time of drug exposure; see text for details. **c** Data (circles) is fit to a spontaneous mutagenesis-only model; values in inset indicate maximum-likelihood estimates of the mutation rate *µ*, with solid line showing simulated Luria-Delbrück distributions. **d** As in c, but for a model with both spontaneous and induced mutagenesis, with rates *µ* and *µ*ind, respectively.

To test this, we devised a compound selection – termed *canUVanine* – that mimics drug-induced resistance by pairing transient exposure to ultraviolet light (UV), a common environmental mutagen, with selection on the drug canavanine. In effect, it emulates a selective agent that both induces resistance and then selects for it, introducing a controlled dose of stress-induced mutagenesis immediately prior to selection. We grew three populations of barcoded yeast to saturation and selected for mutants by passaging into canavanine-containing media, but just prior to selection two of the populations were transiently exposed to UV to complete the canUVanine selection. As before, mutants were grown to saturation in selective media, passaged to ensure barcode stability, and sequenced before and after selection (Fig. 4b). Mutant barcode counts were combined to equalise pre-selection population sizes across all three populations, and the calibration from reads to absolute mutant counts was fixed by fitting the non-UV-exposed population to the canonical Luria-Delbrück distribution; see SI for experiment design and analysis details.

We then fit all three populations to a model with only spontaneous mutations using a maximum likelihood approach. The population that was not exposed to UV was found to fit well, with an inferred spontaneous mutation rate of *µ ∼* 3 × 10^−7^ – comparable with our prior experiment (Fig. 3) and published values [14, 47]. However, the best-fit Luria-Delbrück distributions (both with *µ* ∼ 1 × 10^−6^) for the two UV-exposed populations failed to match the experimentally-measured distributions, Fig. 4c.

We then modelled induced mutagenesis as the probability *µ*ind that the selective environment induces resistance in a previously wildtype cell. The data was once again fit by maximum likelihood, this time simultaneously inferring best-fit values of both the spontaneous rate *µ* and the induced rate *µ*ind independently for each population, Fig. 4d. The population that was not exposed to UV was once again fit best by a spontaneous mutation rate *µ* ∼ 3 × 10^−7^ and an induced mutation rate *µ*ind statistically indistinguishable from 0 (SI Fig. S6b) – indicating no support for induced mutagenesis in the population not exposed to UV.

In contrast, maximum likelihood found both spontaneous (*µ* ∼ 2.2 × 10^−7^ and 2.6 × 10^−7^ in populations 1 and 2, respectively) and induced mutagenesis (*µ*ind ∼ 1.9 × 10^−6^ and 1.5 × 10^−6^ in populations 1 and 2, respectively) in the UV-exposed populations. Notably, the inferred rate of spontaneous mutagenesis *µ* was now comparable to that of the no-UV population, while the inferred *µ*ind agreed well with independent measurements from mutant colony counts before and after UV (*µ*ind ∼ 2.2 × 10^−6^ and *µ*ind ∼ 1.5 × 10^−6^ for UV populations 1 and 2, see SI for details). Plotting the inferred distribution alongside the experimental data in Fig. 4d, we observe a good fit – indicating the presence of induced mutagenesis in the canUVanine-selected populations and validating the power of BarFly in quantifying the relative contributions of spontaneous and induced mutations to the evolution of drug resistance.

### E. BarFly as a multiplexed mutation-rate assay

Besides its historic use in identifying the mechanism of adaptation [10], Luria-Delbrück assays are also commonly used simply to estimate mutation rates in microbes and cell lines. In this mode, one typically needs only ≤ 100 replicates to get a good estimate of the mutation rate [15]. We reasoned that, with its enhanced throughput, BarFly could be used to measure thousands of mutation rates in a single experiment, by splitting up the ‘budget’ of, e.g., 10^5^ barcodes across multiple strains, allowing us to measure the mutation rates of 10^3^ different strains with 100 replicates each, Fig. 5a.

**FIG. 5.**
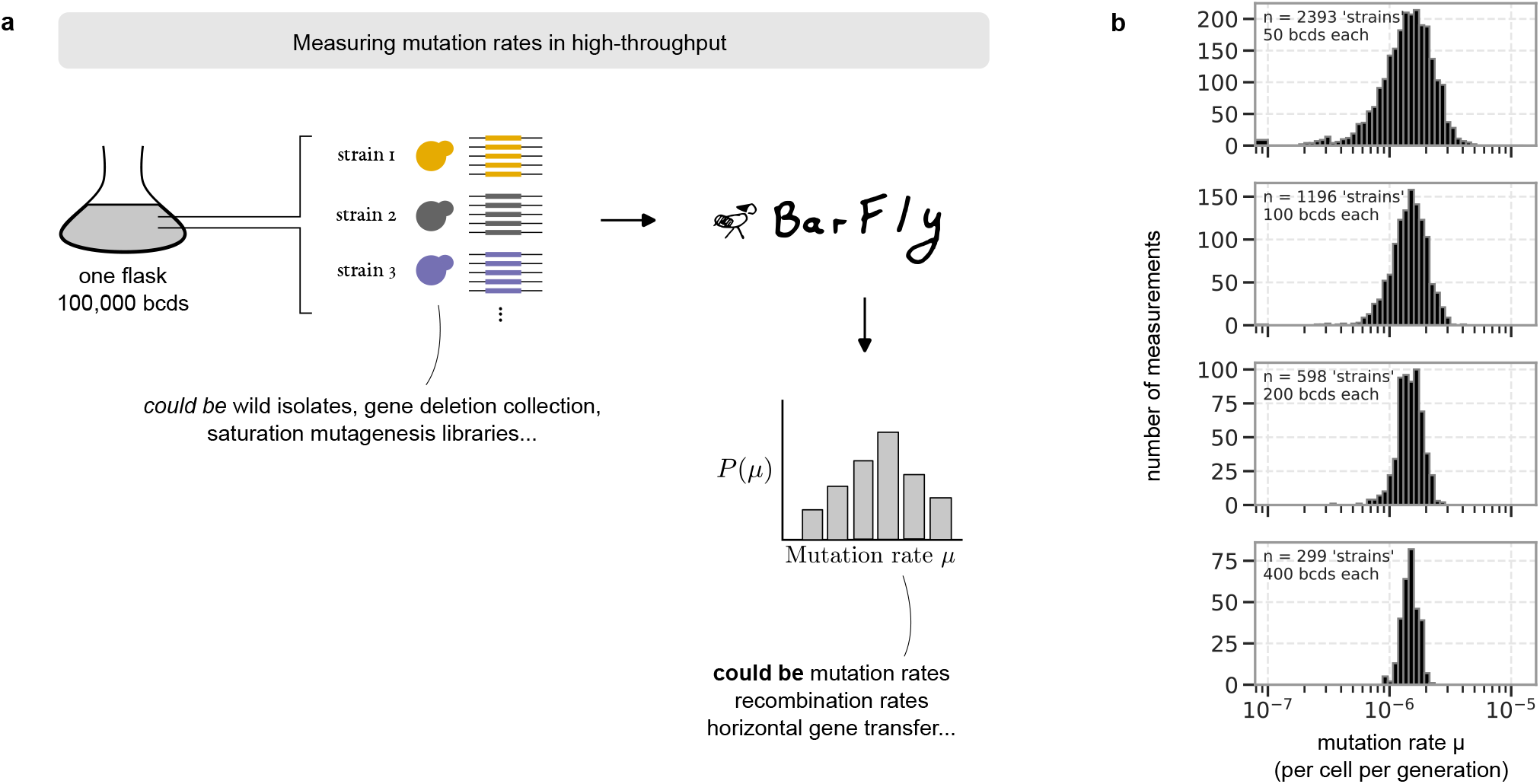
BarFly outperforms traditional fluctuation assays. **(a)** A schematic of BarFly used to perform a *deep mutation rate scan* (D*µ*S) to make a multiplexed measurement of mutation rates or other similar evolutionary rates. **(b)** We randomly assigned barcodes from Fig. 2 to virtual ‘strains’ to simulate a D*µ*S. Mutation rates for each strain were estimated by maximum likelihood; figure shows histograms of inferred mutation rates for strains with a varying number of barcodes each.

How many barcodes per strain are required for a reliable estimate? We split the barcodes from Fig. 2 to generate thousands of datasets at a varying number of barcodes per virtual ‘strain’ and inferred the mutation rate of each ‘strain’ by maximum likelihood. The distribution of inferred *µ* is sharply peaked (< 3-fold error) even with as few as 50 barcodes per strain (Fig. 5b), with a median *µ* ∼ 1.5 × 10^−6^ in good agreement with the value from fitting the full distribution with 10^5^ replicates. We conclude that BarFly could perform a highly-multiplexed, one-pot mutation rate assay with throughput beyond the current state-of-the-art [16, 31], with further improvements in, e.g., barcodes per experiment increasing throughput and/or accuracy.

## III. DISCUSSION

We have described BarFly: a high-resolution Luria-Delbrück assay in which DNA barcodes are used to sample the long-tailed statistics of adaptation to a new environment. DNA barcodes are often used to track lineages in development and evolution [40, 41]; here, we adapt it for the conceptually different purpose of repeatedly sampling from a broad distribution. By going beyond summary statistics to capture the shape of the Luria-Delbrück distribution, BarFly can determine how an evolving population generates heritable variation to adapt. We found that mismatch repair mutants (Fig. 3) only increased the spontaneous mutation rate *µ*, whereas transient UV exposure (Fig. 4) left *µ* unchanged but added an additional mode of adaptation in the form of induced mutations.

We anticipate that BarFly will be broadly applicable to identifying modes of drug resistance in cancers and pathogens. A key strength of this phenomenological approach is that it does not rely on specific hypotheses about the underlying molecular mechanisms; instead, the assay provides a starting point for follow-up mechanistic studies. For example, BarFly could determine whether resistance to an uncharacterised chemotherapy drug arises from spontaneous genetic mutations [10], drug-induced mutations [2, 57], non-genetic mechanisms [3, 5–7, 25, 26], or a combination of the above. Follow-up work could then determine the underlying molecular mechanisms – potentially by comparing BarFly results before and after a genetic perturbation.

More generally, the development of BarFly motivates revisiting the theory of the Luria-Delbrück process. Although there is a long history of incorporating different biological processes into calculations of the Luria-Delbrück distribution [24], these models have not typically been developed in the context of inferring multiple underlying mechanisms from a measured distribution. Consequently, we do not yet know which mechanisms are, in principle, identifiable from the Luria-Delbrück distribution. For example, both mutant fitness [36] and phenotypic switching [25] are predicted to alter the tail of the distribution, but whether they produce distinguishable statistical signatures remains an important open question.

Several technical aspects of the assay could be improved. Fitting an empirical distribution requires estimating the absolute number of mutant cells from read counts; here we obtain an initial estimate by colony counting and refine it during fitting, but future implementations could include a spike-in of barcoded cells at known abundance as an internal calibration standard [58, 59]. A further confounding factor is variation in mutant fitness, which causes competition between lineages and distorts barcode frequencies away from the Luria-Delbrück distribution. We found these frequencies remained stable for canavanine-resistant mutants, indicating no detectable fitness differences, but this may not hold in other selective environments and would require additional calibration.

An orthogonal application of BarFly is as a multiplexed mutation rate assay (Fig. 5). Quantifying mutation rates remains important across biology, evidenced by recent efforts to do so in high-throughput [16, 22]. Notably, the Luria-Delbrück assay has recently been adapted to measure other rates, such as those of horizontal gene transfer [19] or plasmid loss [60], suggesting that BarFly could quantify diverse evolutionary rates in high-throughput.

In conclusion, we have argued that the Luria-Delbrück approach remains relevant to modern problems in biology. While our experimental methods are possible only because of recent technological advances, the conceptual basis of this work remains that of Luria and Delbrück: to use the statistics of adaptation to study the ebb and flow of heritable variation within a population.

## CODE AND DATA AVAILABILITY

Sequencing reads have been deposited at the Sequence Read Archive under the NCBI BioProject PRJNA1537973. Analysis code is available at github.com/kabirhusain/dnabarcodedluriadelbruck.

## ACKNOWLEDGMENTS

We thank Riccardo Ravasio, Rama Ranganathan, Craig DeValk, Rathi Kannan, and the members of the Pincus and Murugan labs for scientific and technical discussions. We thank Jake Cornwall-Scoones, Harry-Luke McClelland, Marco Cosentino Lagomarsino, Archishman Raju, and Riccardo Ravasio for a critical reading of the manuscript. The authors acknowledge the use of the UCL Myriad High Performance Computing Facility (Myriad@UCL) and the DIAS cluster in the Department of Physics and Astronomy, and associated support services, in the completion of this work. KH thanks Victoria L for helping move this work forward. This work was supported in part by National Institutes of Health grant RM1 GM153533 (to D.P.). AM acknowledges support from the National Science Foundation through PHY-2310781, the Center for Living Systems (grant no. 2317138) and from NIGMS of the National Institutes of Health under award number R35GM151211.

