## Supplementary Information for "A DNA-barcoded Luria-Delbrück assay resolves mechanisms of adaptation"

### ※ Mathematical and computational analysis of the Luria-Delbrück distribution

The Luria-Delbrück experiment is simple to describe. Into a flask is inoculated a single wildtype cell, which grows exponentially to a population of many (typically $> 10^5$) cells. In this period of exponential growth, 'mutants' arise. Mutants also grow exponentially, though they may not have the same growth rate as the wildtype - in general, their growth rate is sampled from a distribution of fitness effects (DFE) that includes deleterious, neutral, and potentially beneficial mutations.

The entire population is then *selected* by transitioning to an environment in which the wildtype genotype cannot grow. However, a fraction of the mutants that arose are able to survive in this environment. The Luria-Delbrück distribution is the distribution of the number of these adaptive mutants in replicate experiments.

The reason to study this distribution is that its quantitative form depends sensitively on details of the mutational process. By measuring the distribution, and fitting it appropriately, the parameters that governs the dynamics of standing genetic variation can be, in principle, determined. Famously, Luria and Delbrück measured only the first two moments of the distribution and from that inferred the existence of pre-existing genetic variation.

#### 1.1 Notation

We will consider for simplicity (unless otherwise noted) that the population starts with a single wildtype cell and grows exponentially with growth-rate $\lambda_0$ to a final population of size $N$. For now, we consider only spontaneous genetic mutations – with other mechanisms of adaptation (e.g., transient phenotypic heterogeneity, or induced mutations) considered later. Mutations arise with a rate $\mu_0$ substitutions per base per generation (sbp). Typical numbers for $\mu_0$ are between $10^{-10}$ sbp and $10^{-7}$ sbp for microbial mutation rates in the genome. When mutants arise, they also grow exponentially with growth-rate $\lambda$. It will be useful to consider the ratio of mutant and wildtype growth-rates,

$$\sigma \equiv \frac{\lambda}{\lambda_0}. \tag{1}$$

If $\sigma > 1$ ($< 1$) the mutant grows faster (slower) than the wildtype: $\sigma$ is essentially the fitness of the mutation. We use $\sigma$ rather than the more traditional selection co-efficient $s$ out of convenience; the two are equivalent, with $s = (\sigma - 1)\ln 2$.

Different mutations will, in general, have different effects on fitness. This is traditionally captured by the distribution of fitness effects (DFE). In principle, the mutation rate $\mu_0$ and the DFE completely specify the dynamics of genetic variation.

How does these parameter enter the Luria-Delbrück distribution? Recall that the Luria-Delbrück experiment measures the number of mutants $m$ that are adaptive (i.e. survive) in the new environment. It should be emphasised that the Luria-Delbrück distribution depends *only* on the properties of these adaptive mutations. That is, only a subset of the evolutionary parameters parameterise the distribution.

In particular, the 'mutation rate' that goes into the Luria-Delbrück distribution is not the bare mutation rate per base pair $\mu_0$, but instead the rate $\mu$ at which adaptive mutations arise. This is often written in terms of the per-base-pair rate $\mu_0$ and the number of adaptive mutations available to the organism (i.e. the 'mutational cross-section') $C$,

$$\mu = \mu_0 \times C, \tag{2}$$

which is often referred to as the *phenotypic mutation rate* in the Luria-Delbrück literature. For the typical "fluctuation assays" done in the lab (e.g., 5FOA or canavanine selection in yeast), the target is a loss of function mutation in one or more genes, with $C \sim \mathcal{O}(10^2)$ (estimated to be 236 basepairs for canavanine in [1]) and $\mu$ correspondingly in the range of $10^{-8}$ to $10^{-6}$.

#### 1.2 Basic properties of the distribution: $P_0$ and the power-law tail

It is easy to characterise the extremes of the Luria-Delbrück distribution – at $m = 0$ (no mutants observed) and $m \gg 1$ (many mutants seen – the famous jackpots). For the former, note simply that if we see no mutants, then no *mutations* have occurred. The number of mutations that occur is Poisson distributed with mean $\mu N$, and so:

$$P(m = 0) = e^{-\mu N} \tag{3}$$

This is often referred to as the "$P_0$" value, and it has been used (even in the original Luria-Delbrück paper [2]) to estimate mutation rates from a fluctuation assay [3].

We can also directly compute how the tail of the distribution decays, at least for the case where all adaptive mutations have a single fitness $\sigma$. To do so, suppose that the Luria-Delbrück experiment starts at time $t = 0$ with one cell, and ends at time $t = T$ with $N = e^{\lambda_0 T}$ cells.

The tail of the distribution is dominated by rare mutations that occur early, and so we consider only a single mutation that occurs at some time $0 < \tau < T$, and consequently gives rise to a mutant lineage of at least $m$ mutants, where $m = e^{\lambda(T-\tau)}$. At time $\tau$, the population size is $n(\tau) = e^{\lambda_0 \tau}$, and so the probability that a mutation occurred before time $\tau$ is

$$1 - e^{-\mu n(\tau)} \approx \mu e^{\lambda_0 \tau},$$

which is, equivalently, the probability that at least $m$ mutants were observed. Now, express $\tau$ in terms of $m$:

$$\lambda(T - \tau) = \ln m \Rightarrow \tau = T - \frac{1}{\lambda} \ln m$$

and we obtain

$$\begin{aligned} \text{Prob}\,(\#\text{ mutants } > m) &\approx \mu N \exp\left(-\frac{\lambda_0}{\lambda} \ln m\right) \\ &= \frac{\mu N}{m^{\lambda_0/\lambda}} \\ &= \mu N\, m^{-1/\sigma}, \end{aligned} \tag{4}$$

which is the famous power-law tail of the Luria-Delbrück distribution. This long tail is simply the mathematical expression of the classic 'jackpot' explanation for the Luria-Delbrück process. Naturally, the power-law tail must be eventually truncated, as the number of mutants $m$ cannot exceed the total population size $N$. This will emerge from the more detailed calculation below.

#### 1.3 Derivation of the cumulant generating function

We follow here the treatment of [4]. Consider a population that starts with $n_0$ wildtype cells and grows to $N$ cells. Mutations occur with probability $\mu \ll 1$ at each cell division. Define the indicator function $X_n$, which tracks whether a mutation has occurred as the population transitions from $n$ cells to $n + 1$ cells:

$$X_n = \begin{cases} 0 \text{ with probability } 1 - \mu \\ 1 \text{ with probability } \mu \end{cases}$$

Each mutation gives rise to a mutant lineage of size $Y_n$,

$$Y_n = \left(\frac{N}{n}\right)^{\sigma} X_n$$

where $\sigma$ denotes the ratio between mutant and wildtype growth rates. The total number of mutants produced is simply:

$$m = \sum_{n=n_0}^{N} Y_n$$

We will compute the log of the Laplace transform of $P(m)$, also known (up-to the sign of the argument) as the cumulant generating function:

$$\begin{aligned}
\ln \tilde{P}(s) &\equiv \ln \int_0^\infty dm\, e^{-sm} P(m) = \ln\langle e^{-sm}\rangle \\
&= \sum_{n=n_0}^{n=N} \ln \left\langle \exp\left(-s\left(\frac{N}{n}\right)^\sigma X_n\right)\right\rangle \\
&= \sum_{n=n_0}^{n=N} \ln\left(1-\mu+\mu e^{-s(N/n)^\sigma}\right) \qquad (5)
\end{aligned}$$

where in the last line we have used the definition of $X_n$. We now make a change of variables to $x = n/N$ and approximate the sum by an integral,

$$\ln \tilde{P}(s) \approx N \int_{x_0}^{1} dx\, \ln\left(1-\mu+\mu e^{-s/x^\sigma}\right) \qquad (6)$$

where $x_0 = n_0/N$.

As $\mu \ll 1$, and if we restrict ourselves to $\mathrm{Re}\, s > 0$, we can expand the logarithm to first order in $\mu$ and perform the integral,

$$\begin{aligned}
\ln \tilde{P}(s) &\approx \mu N \int_{x_0}^{1} dx \left(e^{-s/x^\sigma} - 1\right) \\
&= -\mu N\,(1-x_0) + \mu N \int_{x_0}^{1} dx\, e^{-s/x^\sigma} \\
&= -\mu N\,(1-x_0) + \frac{\mu N}{\sigma} \mathrm{E}_{1+1/\sigma}(s) - \frac{\mu N}{\sigma} x_0\, \mathrm{E}_{1+1/\sigma}\left(s\, x_0^{-\sigma}\right) \qquad (7)
\end{aligned}$$

where $E_p(z)$ is the special function known as the generalised exponential integral, defined as:

$$E_p(z) = \frac{1}{z^{p-1}} \int_z^\infty dt\, \frac{e^{-t}}{t^p}$$

Eq. 7 is, in principle, the end of the calculation. Note, however, that the first and third terms depend on $x_0$, which is of $\mathcal{O}(1/N)$; we simplify the expression by approximating $1 - x_0 \approx 1$ in the first term. The third term remains small until $s < x_0^\sigma \sim N^{-\sigma}$. Interpreting $s$ as the "frequency" corresponding to the mutant number $m$, this means that the term is only in play when $m$ is of order $N^\sigma$. Since we are never, in practice, dealing with $m \sim N$, we neglect that term to arrive at a compact expression,

$$\ln \tilde{P}(s) \approx -\mu N \left(1 - \frac{1}{\sigma} \mathrm{E}_{1+1/\sigma}(s)\right). \qquad (8)$$

The parameters $\mu$ and $N$ only enter as the product $\mu N$, which itself only enters as an over-all scale factor.

#### 1.4 Stability of the distribution under addition

Note that Eq. 8 implies that the Luria-Delbrück distribution is (approximately, see below) stable under addition. Recall that a distribution $P(m)$ is stable if the sum $M = m_1 + m_2$, where $m_1$ and $m_2$ are both sampled from $P(m)$, has the same distribution as $P(m)$ up-to changes in location and scale parameters. This follows directly by realising that the distribution of $M$, $Q(M)$, is obtained by convolution of $P(m)$ with itself:

$$Q(M) = \int_0^M dm_1\, P(m_1) \times P(M - m_1)$$

However, convolutions are simply products in Laplace space, $\tilde{Q}(s) = \tilde{P}(s) \times \tilde{P}(s)$; taking the logarithm of both sides:

$$\ln \tilde{Q}(s) = \ln \tilde{P}(s) + \ln \tilde{P}(s) = -2\mu N \left(1 - \frac{1}{\sigma}\, \mathrm{E}_{1+1/\sigma}(s)\right)$$

which is identical to Eq. 8 except for the substitution $\mu N \to 2\mu N$. Similarly, taking two (or more) samples with different final population sizes $N_1$ and $N_2$ leads to a Luria-Delbrück distribution with the new scale parameter $\mu(N_1+N_2)$. This seemingly minor point is useful in analysing BarFly data, as barcode abundances in a library are rarely uniform and consequently the population size per barcode can vary over an order of magnitude. Combining barcodes to equalise population size prior to selection allows for very good agreement with the theoretical Luria-Delbrück distribution.

It is important to emphasise that the stability of the distribution under addition is only approximate, as the arguments only holds for the approximate form Eq. 8 rather than the full expression Eq. 7. However, as the two expressions differ only when the number of mutants is on the order of the population size $N$ (i.e., a mutation occurs in the first few generations of growth), the approximation is valid over the regime relevant for experimental data.

#### 1.5 Numerical calculations of the Luria-Delbrück distribution

Simulations of the Luria-Delbrück process are done with a discrete Gillespie-like algorithm. We suppose that the experiment starts with a single wildtype cell that grows exponentially to $N$ cells in time $T$. That is, taking the number of wildtype cells, $n(t)$, to be a continuous variable, we have $n(t) = e^{\lambda_0 t}$ where $\lambda_0$ is the wildtype growth rate. The final time $T$ and the final population size $N$ are related through $N = e^{\lambda_0 T}$.

The numerical algorithm takes the following steps:

1. Determine the number of mutations that occur, $r$, by sampling from a Poisson distribution with mean $\mu N$
2. For each mutation $i$ $(1 \leq i \leq r)$, determine the time $t_i$ $(0 \leq t_i \leq T)$ at which the mutation occurred by sampling from $P_\mu(t)$ (see below).
3. Calculate $m$ as $\sum_i m_i$, where $m_i = e^{\lambda(T-t_i)}$ and $\lambda$ is the mutant growth rate.

The simulation first samples the number of mutations that occur during growth. Mutations occur stochastically with a rate $\mu$ per generation, and the number of mutations that occur, $r$, is therefore Poisson-distributed with mean $\mu N$. Note that this is not the Luria-Delbrück distribution, which is instead the number of *mutants* descended from these mutations. For each mutation $i$ that occurs at time $t_i$, the number of mutants produced is $m_i = e^{\lambda(T-t_i)}$, where $\lambda$ is the mutant growth rate. $t_i$ is also a random number and needs to be sampled from the appropriate distribution.

We compute the statistics of $t_i$ in the following way. In a slice of time between $t$ and $t + dt$, mutations occur with rate $\mu n(t) dt$. Normalising over the entire time window from $t = 0$ to $t = T$, we obtain the probability density that the mutation occurs at time $t$:

$$P_\mu(t)dt = \frac{\mu n(t)dt}{\int_0^T dt'\,\mu n(t')} = \lambda_0 \frac{e^{\lambda_0 t}}{N-1} \tag{9}$$

which is normalised over $t = 0$ to $t = T$. By a direct application of inverse transform sampling, $P_\mu(t)$ can be sampled from by generating a random number on the unit interval, $x$, and transforming it as $t = (1/\lambda_0)\ln(1+(N-1)x)$. With $t_i$ in hand, the number of mutants that arise from mutation $i$ is simply $m_i = e^{\lambda(T-t_i)}$. In all simulations we take $\lambda_0 = 1$ to set units of time.

### 1.6 Simulating different adaptive mechanisms

In Fig. 1 of the main text we present Luria-Delbrück distributions arising from different adaptive mechanisms described in the literature. In each example, we contrast the simulated distribution with a canonical Luria-Delbrück distribution, which was obtained with mutation rate $\mu = 2.5 \times 10^{-6}$ (comparable to the msh2$\Delta$ strains adapting to canavanine in Figs. 2 and 3 of the main text). All simulations were done with a population size $N = 10^6$.

Drug-induced mutations were simulated by first sampling a mutant number $m_{\text{LD}}$ from a canonical Luria-Delbrück distribution and then adding to it a sample $m_{\text{ind}}$ from a Poisson distribution with mean $\mu_{\text{ind}}N$ – obtaining a final mutant number $m = m_{\text{LD}} + m_{\text{ind}}$. We used $\mu_{\text{ind}} = 5 \times 10^{-6}$, comparable to rates determined in Fig. 4 of the main text. We used a reduced spontaneous mutation rate $\mu = 10^{-6}$ to keep the overall rate that mutations occur roughly comparable to the canonical distribution presented in the same plot.

A phenotypic delay was implemented by modifying the numerical algorithm above to only consider mutations that occur between $t = 0$ and $t = T - t_{\text{delay}}$, where $T$ is the time at which the selection is applied, and $t_{\text{delay}}$ is the timescale of phenotypic delay (i.e. the time before which a mutant does not express its phenotype). We used $t_{\text{delay}} = 2.5$ generations, similar to estimates suggesting a $\sim 3$ generation delay from experiments in bacteria [5]. We increased $\mu$ to $7 \times 10^{-6}$ to keep the overall number of mutations approximately constant.

To simulate the effects of adaptation occurring via reversible phenotypic switching rather than permanent genetic changes, we suppose that the 'mutation rate' $\mu$ now describes the rate at which cells transition into a transient phenotypic state that – for simplicity – we continue to call mutants. As before, the 'mutants' grow at rate $\lambda$. Unlike genetic mutants, however, these revert back to the wildtype with a rate $\nu = 1/\tau$, where $\tau$ is the timescale over which the phenotypic state is inherited.

We distinguish two cases:

**1. $\lambda < \nu$:** Here, reversion is faster than growth, and new mutants do not have time to establish a lineage. Consequently, the resulting distribution will be Poisson.

**2. $\lambda > \nu$:** If reversion is slower than growth, then the mutant grows out into a lineage but some fraction of its descendants revert back to the wildtype. The simplest deterministic dynamics for the size of the mutant lineages is

$$\frac{dm}{dt} = \lambda m - \nu m,$$

from which we see that the net growth rate of the lineage is simply $\lambda - \nu$. As a result, as far as the dynamics of lineage size go, the reversible phenotypic change is equivalent to an *irreversible* genetic mutation with a renormalised $\sigma$,

$$\sigma_{\text{effective}} = \frac{\lambda - \nu}{\lambda_0},$$

We therefore simulate a Luria-Delbrück distribution with this renormalised $\sigma$. We choose $\mu = 2.5 \times 10^{-6}$, $\lambda = \lambda_0$ (i.e. no intrinsic fitness cost to the mutation), and $\nu = 0.3$ – corresponding to approximately a 3 generation memory for a phenotypic resistant state, consistent with recent inference from experimental data [6].

Note that this argument neglects many sources of stochasticity – in particular, of the reversion process. That (and other stochastic effects) could fundamentally alter the argument of equivalence between mutant fitness $\sigma$ and reversion rate $\nu$; we leave it to future mathematical work to investigate the issue.

### ※ Strain & plasmid construction and handling

#### 2.1 Media and growth conditions

Yeast were grown either in YPD (1% yeast extract, 2% peptone, 2% dextrose) or in synthetic complete (SC) media (1.7 g/L yeast nitrogen base, 5 g/L ammonium sulfate, 36 mg/L myo-inositol) supplemented with the appropriate amino acid drop-out mix. Solid media included 2% agar, while all canavanine selections were done at a final concentration of 100 mg/L of the drug. All growth in liquid media was done at 30C with shaking, with any culture above 10mL being grown in a baffled flask. In the experiments presented in Figs. 2 and 3 of the main text, the initial selection in canavanine was done in semi-solid media containing 0.35% SeaPrep Agarose.

Unless otherwise mentioned, all plasmid cloning was done with XL1-Blue competent cells prepared using the Zymo Mix & Go kit (Zymo Research Cat. # T3002). CRISPR/Cas9 knockouts were performed using a Cas9 plasmid (Addgene # 60847 [7], into which a ura3 or trp1 selectable marker was cloned alongside the existing KanMX cassette, for flexibility of selection).

Guide sequences were cloned into the Cas9 plasmid by blunt end ligation. Briefly, the entire plasmid was linearised in a PCR by primers introducing the desired sequence on their overhangs. 1uL of the PCR product was then combined with 1uL each of DpnI (NEB Cat. # R0176S), T4 PNK (NEB Cat. # M0201S), and T4 ligase (NEB Cat. # M0202S) in a 10uL reaction containing 1x NEB CutSmart buffer (NEB Cat. # B6004S); the reaction was incubated at room temperature for one hour, and then transformed directly into competent cells. Plasmids were miniprepped and verified by whole-plasmid sequencing (Plasmidsaurus).

#### 2.2 Gene deletions

The mismatch repair genes msh2 and msh6 were deleted using CRISPR/Cas9 to cleanly delete the ORFs of each gene. In brief, we cloned guide sequences (`GAAGTTGTCTCTGTGCCAGG` for msh6, `CAAACATAACTTCAGCAGAG` for msh2) into the Cas9 plasmid, and made 100bp double-stranded repair templates (containing 50bp of homology upstream and downstream of the desired ORF) by PCR. Each knockout was done with 500ng to 1ug of pCas plasmid and 1 to 5ug of repair template transformed into yeast by a standard Lithium acetate protocol [8]; knockouts were verified by diagnostic PCR.

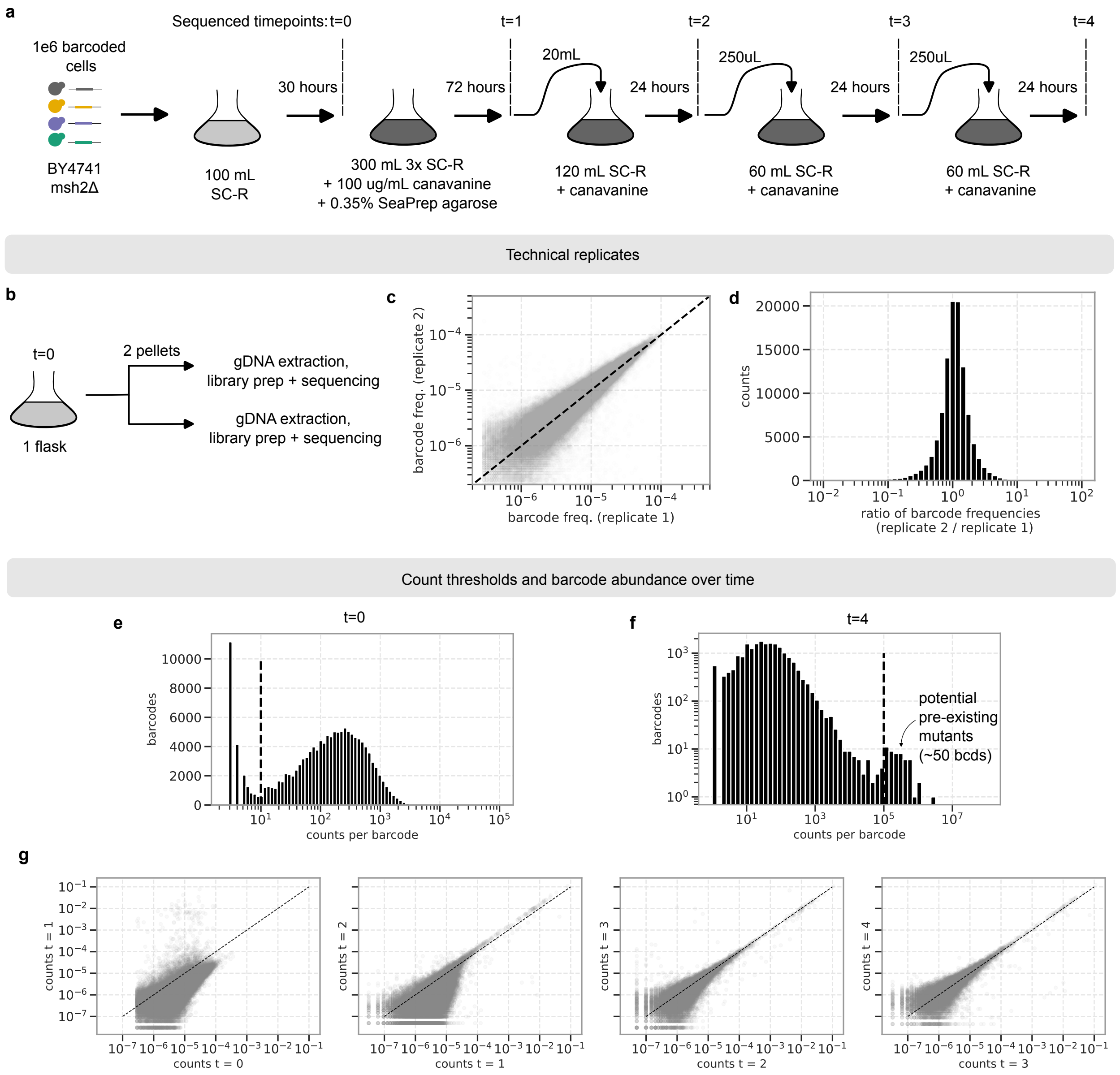

Figure S1: **Experimental details and initial analysis for data in Fig. 2 of the main text.** **a** Outline of the experiment. **b** To assess technical noise, we sequenced barcodes from libraries prepared independently from two samples of the same flask (at the initial timepoint $t = 0$). **c** Scatter plot of barcode frequencies, and **d** ratio of barcode frequencies between technical replicates. **e** Barcode frequencies at the initial and **f** final timepoints. Vertical dashed lines show thresholds used to exclude sequencing noise and lineages that were likely to harbour pre-existing mutants, respectively. **g** Comparison of barcode frequencies at subsequent timepoints. Dashed lines in (c) and (g) are the $x = y$ line. Time-points $t = 2$, $t = 3$, and $t = 4$ are separated by approximately 16 generation of growth ($\log_2(60/.250) \approx 7.91$).

#### 2.3 Barcoding

To genomically barcode yeast, we developed a high-efficiency genomic integration strategy that exploits co-transformation with a linearised Cas9 cassette. Our barcoding system is built around the yeast integrating plasmid pNH604, which integrates into the Trp1 locus by homologous recombination, and carries a Trp1 gene from *Candida glabrata* for selection of transformants. We engineered a barcoding library by introducing a small, 12 nt barcode in the integration cassette region of pNH604 (method below). To increase transformation efficiency, we co-transformed with a linearised Cas9 plasmid targeting the integration region in the genome.

Barcodes were introduced into the yeast integrating plasmid pNH604 by PCR. The entire plasmid was amplified with primers that bring in a 12nt barcode (in blocks of 4, separated by two As to avoid accidentally introducing an unwanted restriction enzyme site: `NNNNAANNNNAANNNN`) along with BamHI sites on the terminal ends of the PCR fragment. The fragment was treated with DpnI, digested with BamHI, self-ligated with T4 ligase, and transformed into homemade electrocompetent XL1-Blue *E. coli* to obtain an estimated $10^6$ transformants. After overnight growth in LB at 37C, the plasmid library was miniprepped and frozen.

Our genetic background, BY4741, is Trp+ and contains a functional Trp1 gene. To barcode these cells, we used a two-part strategy. First, the functional Trp1 gene was deleted and replaced with a 'landing pad' using a Cas9 guide (`TTGCGGCTTGCAGAGCACAG`) that targets the Trp1 coding region between the homology arms used for integration of pNH604, and supplying the landing pad as a repair template. The landing pad contained a synthetic CRISPR target site `TCCCCAATCGTGGAGTGAAG` (characterised in [9]; not present elsewhere in the yeast genome).

The landing pad was then overwritten with the barcode cassette, aided by co-transformation with a linearised Cas9 targeting the synthetic cut-site in the landing pad. Approximately 10 $\mu g$ of the barcode plasmid library and 5 $\mu g$ of the Cas9 plasmid were digested overnight with PmeI and XhoI, respectively. After enzyme inactivation, the digested plasmids were combined and transformed into yeast using a modified Lithium-Acetate method. In brief, an overnight culture of yeast was diluted $\sim 1 : 10^5$ into 50mL of YPD. The following morning, cells were harvested between OD600 0.5 and 0.7 by centrifugation, washed once with 1mL of LiTe (100 mM Lithium acetate, 10 mM Tris pH 8, 1 mM ethylenediaminetetraacetic acid) and resuspended to 500 uL with LiTE in a 15mL conical, to which was added 2.5mL of LiTEPeg, 150 uL of single-stranded salmon sperm, and the digested plasmids. The mixture was incubated at 30C for 25 min, heat shocked at 42C for 40 min, then spun down and resuspended (by gentle inversion) in 5 mL of synthetic complete media lacking tryptophan (SC-W), plating dilutions to estimate transformation efficiency. Typical efficiencies were between 50,000 and 500,000 transformants.

Post-transformations, yeast libraries were grown to saturation in 250mL of SC-W at 30C over two days, passaged once 1:100 into 100 mL of SC-W to dilute untransformed cells, grown to saturation again overnight, and aliquots frozen in YPD supplemented with 15% glycerol.

### ※ BarFly Protocol

#### 3.1 General protocol: growth and selection

All BarFly experiments described here followed a common protocol (described below, including any major variations). Figs. S1, S2, and S3 show a schematic of all experimental outlines, including the precise values of, e.g., the volumes of cultures used (which varied slightly between experiments).

To begin a BarFly assay, a glycerol stock of barcoded cells was thawed, inoculated into 50mL of YPD, and grown overnight at 30C. The next day, we bottlenecked the population to begin the exponential growth phase of the Luria-Delbrück protocol. A small portion (typically 1uL to 10uL, corresponding to $10^5$ or $10^6$ cells) was transferred into approximately 100mL of synthetic complete media lacking arginine (SC-R). We used SC-R at this stage as our initial characterisation found that canavanine addition to cells grown in SC-R resulted in an immediate cessation of growth, while cells grown in YPD underwent a further two or three doublings before stopping. Previous work [1] found that growth of the culture after the addition of the drug distorts the Luria-Delbrück distribution; we chose SC-R to minimise this.

The culture was grown with shaking at 30C for 30 to 48 hours until saturation, and a few (two or three) 1.5mL aliquots were removed, spun down, and the cell pellets frozen at -20 to prepare genomic DNA for barcode sequencing later. A small portion was also plated onto either YPD or SC-R to determine the total number of cells present, $N_{\text{total}}$. In addition, serial dilutions were plated onto SC-R + canavanine to estimate the total number of mutants $M$. Both these measurements were used in the downstream analysis of the BarFly data, see below.

The remainder of the culture was then subject to canavanine exposure to select for mutants. The entire amount was spun down and resuspended in a large volume ($> 250mL$) of either SC-R + canavanine with (data in Figs 2, 3 or the main text) or without (data in Fig. 4) 0.35 % SeaPrep agarose. We initially used SeaPrep agarose to allow mutant lineages to develop as spatially-separated colonies in 3D, reasoning that by doing so we would minimise any colony-to-colony variation in, e.g., the number of cells. Later experiments found that selecting in liquid media made no obvious difference: e.g., data in Fig. 3 of the main text and from the no-UV culture in Fig. 4 of the main text were done with the same population of barcoded yeast, and differed only in the use of Seaprep – and yet both fit to the canonical Luria-Delbrück distribution with the same inferred mutation rate $\mu$. To ensure that mutant cells had enough nutrients to grow, we used triple-strength (3x) SC-R at this stage.

Experiments resuspended in SeaPrep-containing media were poured as a thin layer into a baffled Fernbach flask and were left to solidify in an ice-water slurry for approximately three hours. They were then carefully transferred to the shelf of a 30C incubator and grown without shaking. Experiments without Seaprep were grown at 30C with shaking. Note that the shaken cultures clearly did not produce 'colonies' in the sense of spatially-distinct aggregates arising from a single cell. Nonetheless, in what follows, we will continue to use 'mutant colonies' to refer to the lineages established by single mutant cells after the exposure to canavanine.

After 3 days, the cultures were removed from the incubator – those in Seaprep were first homogenised by shaking for two hours. Cells were spun down, and a portion of the culture (amounting to approximately 5%) passaged into a fresh flask of SC-R + canavanine, which was left to grow at 30C with shaking. From the remainder of the culture, we once again froze cell pellets corresponding to approximately 1.5mL of saturated culture, as well as plated dilutions onto SC-R and SC-R + canavanine to count colony forming units. Typically, we found that mutant lineages had expanded $\sim 10^5$-fold, indicating that $> 10^3$ cells from each mutant colony were seeded into the next round of growth.

The next day, cultures were once again removed from the incubator and cell pellets frozen while plating a small portion onto SC-R and SC-R + canavanine to quantify cell counts. To passage the experiment, we diluted a fraction of the culture (see Figs. S1, S2, S3 for experiment-specific values) into fresh SC-R + canavanine. This was repeated once more the following day before the final set of cell pellets were spun down and frozen to prepare for genomic DNA extraction and barcode sequencing.

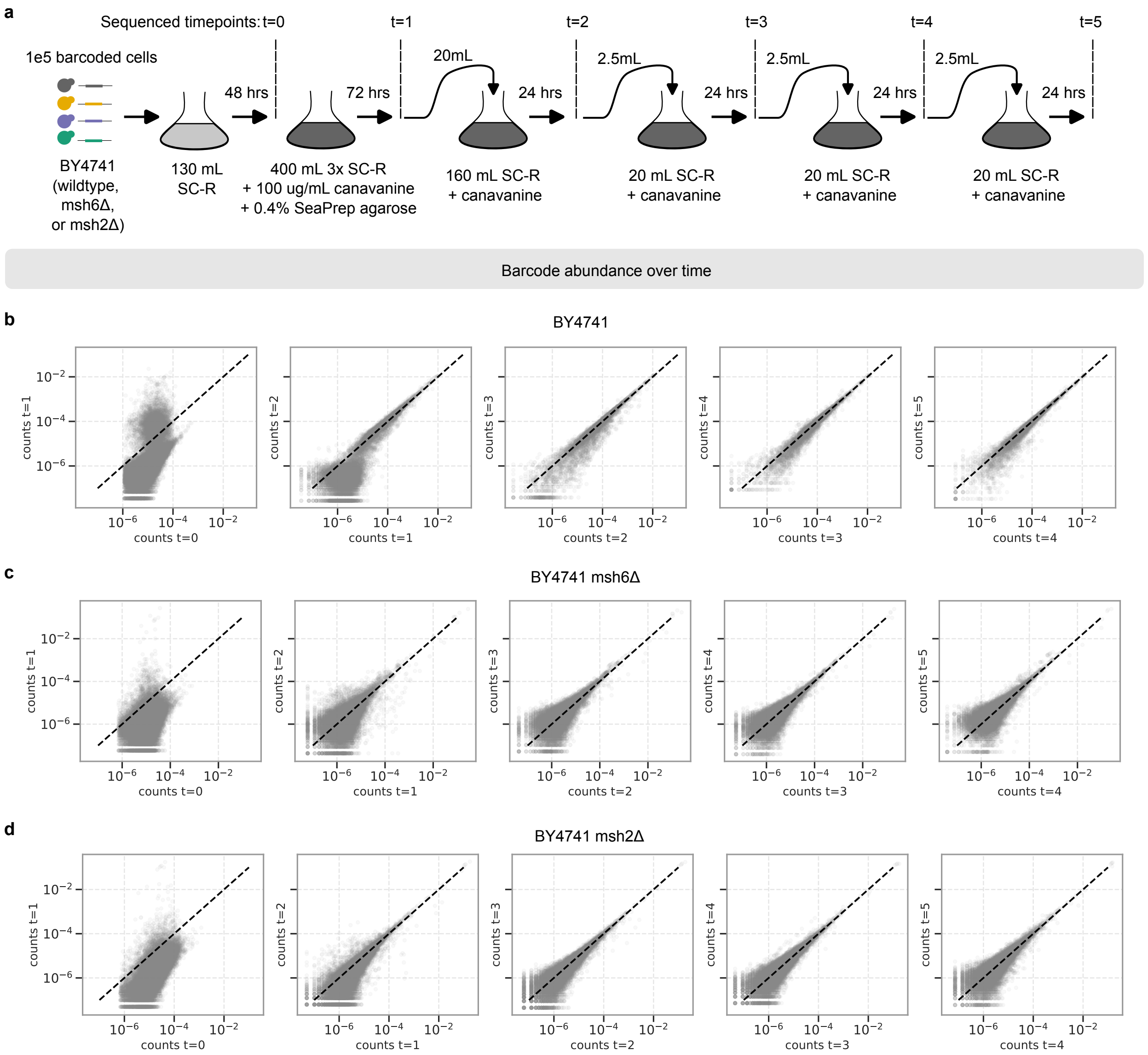

Figure S2: **Experimental details and barcode abundance for data in Fig. 3 of the main text. a** Outline of the experiment. **b, c, d** Comparison of barcode frequencies at subsequent timepoints. Dashed lines are the $x = y$ line.

#### 3.2 UV exposure

Two of the three populations analysed in Fig. 4 of the main text were transiently exposed to ultraviolet light to induce mutagenesis immediately prior to canavanine selection. To do so, flasks containing the saturated cultures were placed under a 302 nm UV overhead light in an Axygen GD-1000 gel imager for 30 minutes. Before exposure, the CFU of each culture was determined via serial dilution onto SC-R plates and by spreading 1mL onto SC-R with canavanine plates. During exposure, the samples were swirled by hand every 5 minutes to avoid settling and to reset the UV light which would turn off at that timescale. Plating onto SC-R and SC-R with canavanine was then repeated and an aliquot for sequencing was taken from each sample. The remainder of the cultures were spun down and resuspended into 1.2L of SC-R with canavanine then placed in a 30 C incubator to continue the BarFly protocol (Fig. S3).

### ※ Barcode Sequencing

#### 4.1 Library preparation and sequencing

Genomic DNA was extracted from cell pellets following a previously described protocol [10]. We prepared amplicon libraries for sequencing by a two-step PCR. In brief, the first round of PCRs consisted of a short (5-cycle) reaction using approximately 10ng to 50ng of genomic DNA as a template. We used one of two primer designs in the reaction, either:

Forward #1: CACTCTTTCCCTACACGACGCTCTTCCGATCT $[N]_n$ GGATCCGATCATGCTT
Reverse #1: TGACTGGAGTTCAGACGTGTGCTCTTCCGATCT $[N]_n$ CAGAAAACGTCATGGAG

or

Forward #2: CACTCTTTCCCTACACGACGCTCTTCCGATCT $[N]_n$ GGATAAAATGTGATAACTAATCAGC
Reverse #2: TGACTGGAGTTCAGACGTGTGCTCTTCCGATCT $[N]_n$ AAACAATCAATGCCAGAG

Each pair anneals to the barcode locus, with the barcode itself positioned to be read within the first few bases of Read 1 on an Illumina sequencer. However, the two primer pairs are designed such that opposite strands of the DNA end up being sequenced, introducing sequence complexity and reducing the amount of, e.g., PhiX or other high-complexity library that needs to be spiked in. The primers also introduce a variable length unique molecular identifier (UMI), denoted $[N]_n$ (with length $n = 8$ to $n = 10$), as well as partial Illumina adaptors (rendered in blue above). The PCRs used NEB Q5 Hot Start DNA polymerase as per the manufacturers instructions, annealing at 55C and extending at 72C for 30s during each of the five cycles. After thermocycling, the PCR product was cleaned with magnetic beads (Aline PCR Clean DX), used at 1.2x following manufacturers instructions, with DNA eluted in 15uL of water. The entirety of the eluted DNA was used as the template for the second round of PCR.

The second round of PCR consisted of 20 cycles, also performed with NEB Q5. We used the following primer design, bringing in the remainder of the Illumina adaptors as well as introducing i5 and i7 indices for identifying samples when multiplexing:

Forward: AATGATACGGCGACCACCGAGATCTACAC [i5 index] ACACTCTTTCCCTACACGAC
Reverse: CAAGCAGAAGACGGCATACGAGAT [i7 index] GTGACTGGAGTTCAGACGTG

The PCR products were once again cleaned with magnetic beads, and a small portion run on a gel for verification. DNA concentration was quantified using the Qubit HS dsDNA dye, and all samples (from, e.g. different timepoints) were combined to obtain an equimolar mix. The pooled library was run out on a gel

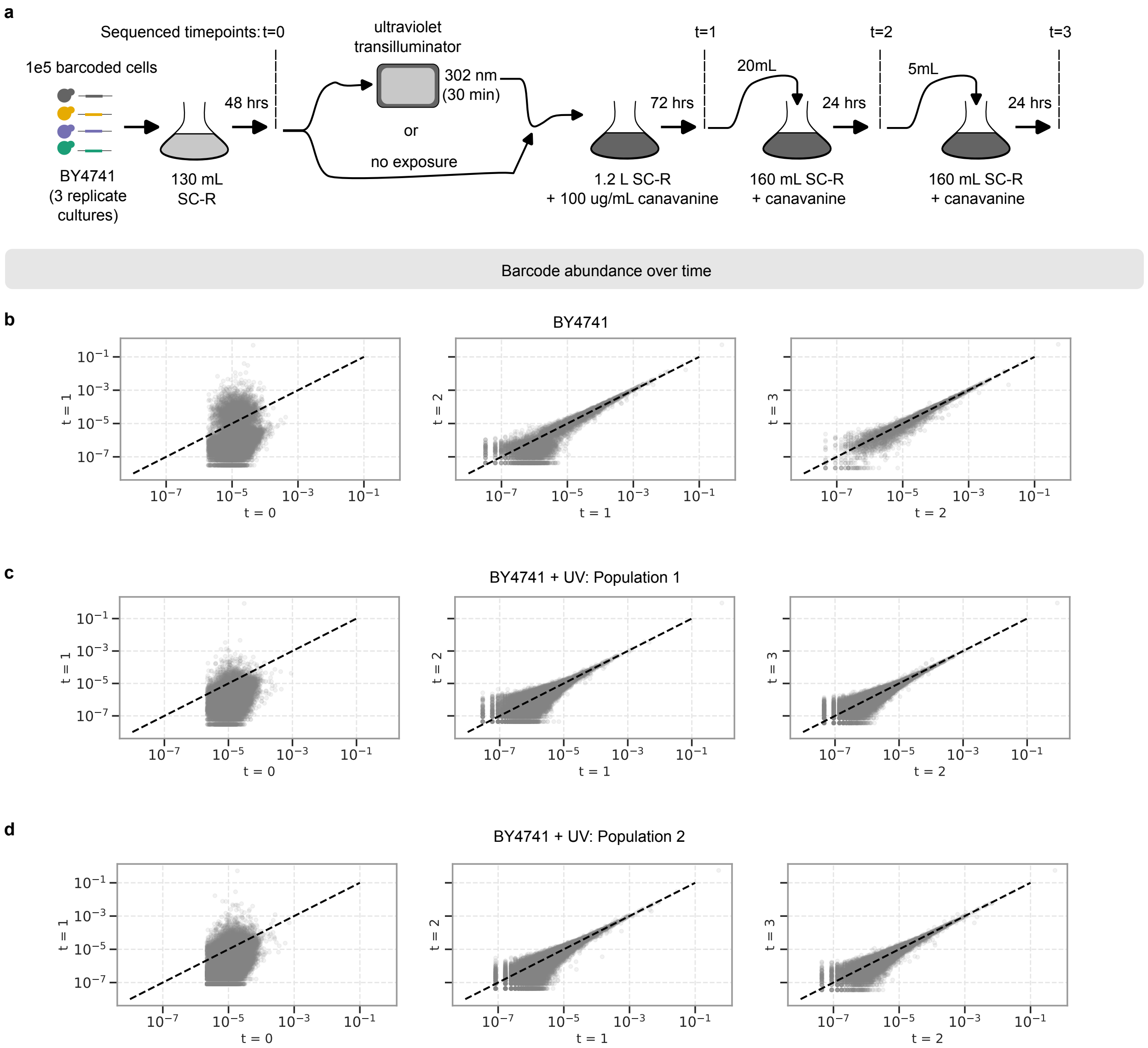

Figure S3: **Experimental details and barcode abundance for data in Fig. 4 of the main text. a** Outline of the experiment. **b, c, d** Comparison of barcode frequencies at subsequent timepoints. Dashed lines are the $x = y$ line.

and the band corresponding to the expected library size cut out and purified with a Zymo silica column using Zymo agarose dissolving buffer according to manufacturers instructions. The final library was quantified and paired-end sequenced on either a NextSeq 500 or a lane of a NovaSeq at the University of Chicago Sequencing Core.

#### 4.2 Barcode extraction and error-correction

Barcodes were identified from sequencing reads by a custom pipeline implemented in Python. Paired-end reads were analysed using regular expressions to identify, in Read 1, a UMI sequence, the primer annealing site, and the barcode, as well as a UMI sequence in Read 2. The UMI from both reads was concatenated to obtain a joint UMI sequence, and the number of reads per barcode and UMI recorded. Next, barcodes were error-corrected by a simple heuristic based on the principle that a sequencing error would produce barcodes of lower read count. First, barcodes were ranked by their abundance into a sorted list, and were analysed in order from the most abundant to the least. For each barcode $b_1$ on the list, all barcodes $b_2$ were identified lower down on the list (i.e., with fewer reads than $b_1$) such that the barcode sequences $b_1$ and $b_2$ differ only by one base (i.e., a Hamming distance of 1). The reads and UMIs belonging to all such barcodes $b_2$ were then reallocated to barcode $b_1$, and barcodes $b_2$ removed from the list to prevent double counting. Finally, the number of unique UMIs per barcode were recorded, and used in all downstream analysis. Note that in what follows, and in the main text, we refer to the number of UMIs as “reads” or “counts” for simplicity.

### ※ Data analysis and fitting

#### 5.1 Identifying barcoded lineages

Not every barcode recovered post-error correction corresponds to a real, well-populated lineage: spurious barcodes arise from residual sequencing and PCR error, and are present at very low abundance. To separate genuine lineages from this background, we examined the histogram of UMIs (“counts”) per barcode at the initial time point ($t = 0$), taken immediately prior to the addition of canavanine. This histogram is clearly bimodal, Fig. S1e: a large mode of low-count barcodes, peaked near singletons, sits alongside a well-separated mode at much higher counts corresponding to the genuinely present lineages. We therefore retained only barcodes falling in the upper mode, imposing a lower cutoff on the $t = 0$ counts (shown as a black dashed line in Fig. S1e).

For the msh2Δ and msh6Δ experiments, the distribution of barcode counts at the final timepoint (e.g., Fig. S1f) had an upper mode of outliers in the histogram of final-time counts, separated from the bulk of the distribution. We attributed this small number of lineages to ones that carry a mutation that pre-dates the experiment, i.e. a canavanine-resistant cell that was already present in the founding population rather than one that arose during the Luria-Delbrück growth window. We chose to exclude them, applying a cutoff for barcodes whose final-time count exceeded an upper cutoff placed between the bulk and this outlier mode. Overall, this upper cutoff removed only a tiny fraction of lineages: in the single-strain experiment of Fig. 2 of the main text, 52 of the 119,724 barcodes passing the lower cutoff (0.04%) were excluded; in the three-strain experiment, 0 of 65,789 wild-type barcodes, 6 of 86,832 msh6Δ barcodes and 6 of 43,528 msh2Δ barcodes ($\lesssim 0.01\%$) were excluded. The retained barcodes constitute the set of genuine lineages used in all downstream analysis.

To assay robustness to technical noise, in the single-strain experiment we sequenced the intial timepoint

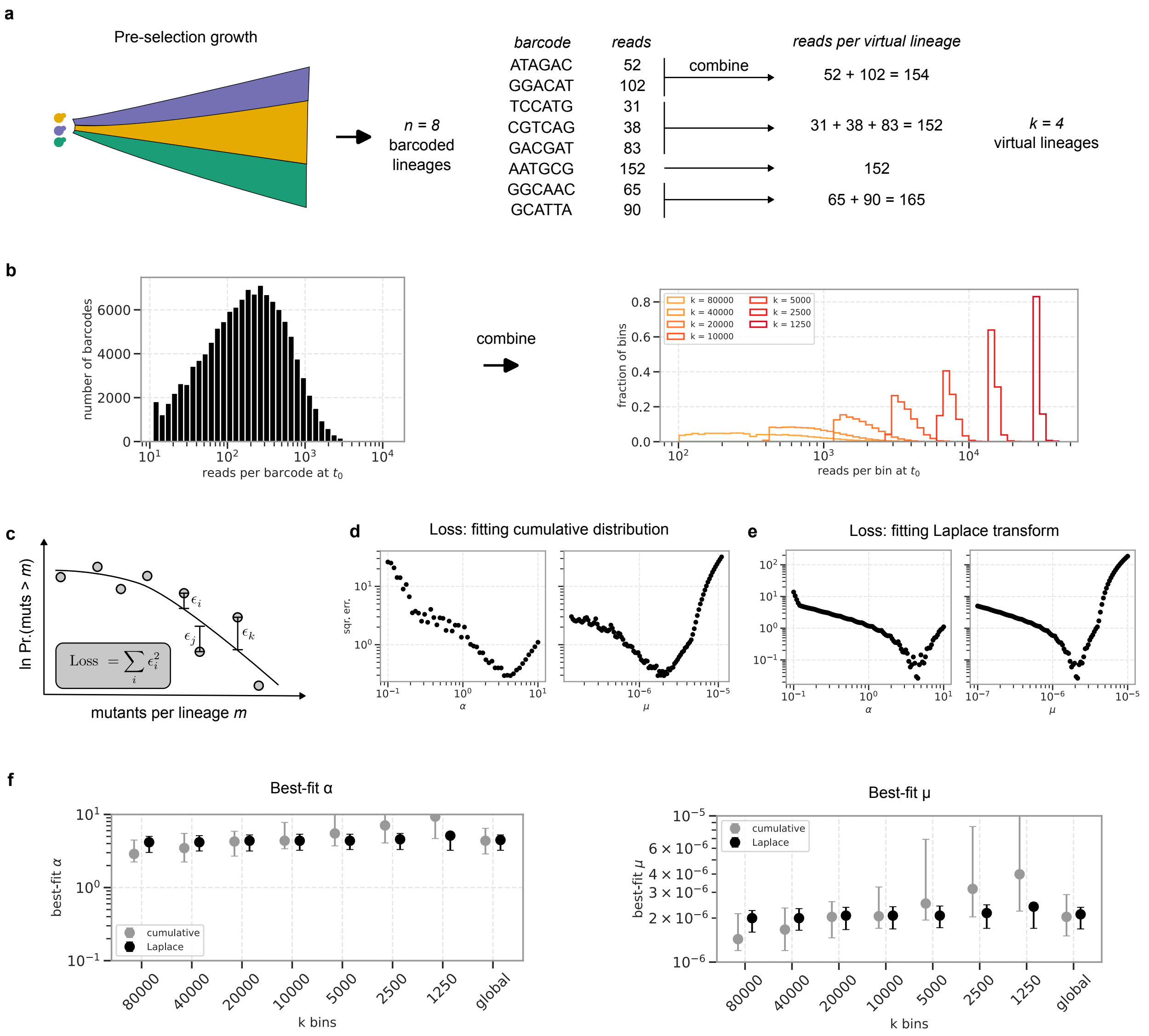

Figure S4: **Combining barcodes and fitting data in Fig. 2 of the main text.** **a** Schematic showing how barcodes can be combined to form 'virtual lineages' with equalised reads pre-selection. **b** (Left) reads per barcode at $t = 0$ (i.e. pre-selection), and (right) reads per virtual lineage after combining barcodes to form $k$ virtual lineages, with varying $k$. **c** Cartoon defining the loss function for least-squares fitting (shown for the complementary cumulative distribution). **d, e** Loss as a function of the fitting parameters – scale factor $\alpha$ and mutation rate $\mu$ – when fitting either the (d) complementary cumulative distribution, or the (e) Laplace transform of the distribution. Plot shows global loss: i.e., summed over $k$. **f** Best-fit $\alpha$ and $\mu$ for different values of $k$.

($t = 0$) twice, preparing two amplicon libraries with independent PCRs from independent genomic DNA extractions. The two technical replicates agree closely (Fig. S1c, d), confirming that the per-barcode counts are broadly reproducible and not dominated by PCR or sequencing noise. As a final quality control, in all experiments we compared the barcode-frequency distribution across successive time points and found it to be stable by the last two time-points (Figs. S1g, S2b, c, d, and S3b, c, d), indicating that there were no appreciable fitness differences between lineages in the selective environment.

#### 5.2 Equalising population size per lineage

Even prior to selection, barcode abundances in the library are far from uniform, and consequently the population size $N$ per lineage varies over more than an order of magnitude. As shown above (*Stability of the distribution under addition*), the Luria-Delbrück distribution is approximately stable under addition. That is, the distribution of $m_1 + m_2 + \ldots$, where $m_i$ is the number of mutants from a population with $N_i$ cells, is approximately the same as a Luria-Delbrück distribution for a population with the same mutation rate, and population size $N_1 + N_2 + \ldots$. We exploited this property to combine lineages into bins of approximately equal founding population size, so that each bin behaves as an independent Luria-Delbrück culture of a common size $N$ and the pooled bins can be compared directly against theory.

Lineages were assigned to bins by a 'longest-processing-time' (LPT) greedy scheduling heuristic, which balances the total $t = 0$ count across bins. Barcodes were sorted in order of decreasing $t = 0$ count and each was sequentially added to the bin with the smallest running total; the resulting bins have more uniform $t = 0$ counts and hence more uniform population sizes. The binning was repeated for a range of bin numbers $k$ (from $k = 1250$ to $k = 80{,}000$ for the single-strain experiment, and from $k = 4000$ to $k = 32{,}000$ for the three-strain experiment), each choice corresponding to an effective per-bin population $N = N_{\text{total}}/k$. For each bin, the total number of mutants was taken to be the sum of the final-time counts of its constituent barcodes.

#### 5.3 Least-squares fit to the Luria-Delbrück distribution

We fit the experimental data to the theoretical Luria-Delbrück distribution by minimising the sum of squared errors between the data and theory. The key parameter is the mutation rate $\mu$, which – in addition to the known pre-selection population size $N$ – completely determines the theoretical distribution. In addition, we included as a fitting parameter a scaling factor $\alpha$ that maps the reads per barcode (or, more generally, a virtual lineage obtained by combining barcodes, see above) at the final time point, $r_i$, to the number of mutant colonies produced by that lineage, $m_i$, as

$$m_i = \alpha \frac{r_i}{\sum_i r_i} M_{\text{cfu}}. \tag{10}$$

Here, $M_{\text{cfu}}$ is an estimate of the total number of mutants produced in the BarFly experiment, obtained by plating a small portion of the experiment onto canavanine plates and counting mutant colonies. As some fraction of viable yeast cells are known to fail to form colonies (a phenomenon broadly known as 'plating efficiency'), we expect $M_{\text{cfu}}$ to underestimate the total number of mutants produced; consequently, while we fit $\alpha$ over a range $\alpha \in [0.1, 10]$, we expect to find $\alpha > 1$. This is true in all of the data-sets analysed, where we consistently recover $\alpha$ between 1 and 5 for all fits.

For each $(\mu, \alpha)$ pair we compared the complementary cumulative distribution function ("CCDF", defined as $P(\text{mutants} > x)$) of the data to a simulated CCDF generated by the discrete simulation algorithm described above (*Numerical calculations of the Luria-Delbrück distribution*). The distributions were evaluated

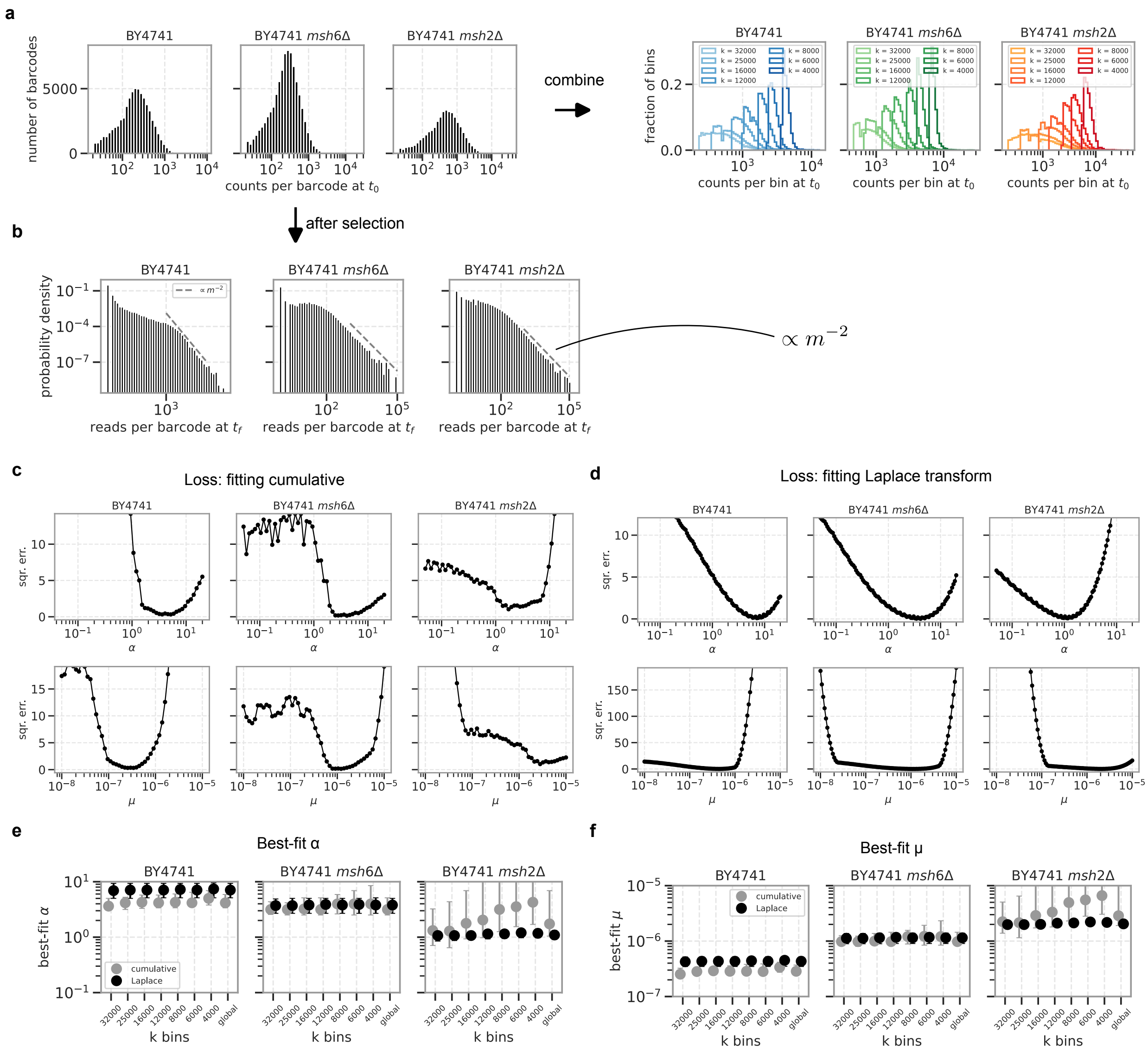

Figure S5: **Combining barcodes and fitting data in Fig. 3 of the main text.** **a** (Left) reads per barcode at $t = 0$ (i.e. pre-selection), and (right) reads per virtual lineage after combining barcodes to form $k$ virtual lineages, with varying $k$. **b** Probability density of reads per barcode at the final timepoint $t = t_f$ (i.e., after selection). Dashed line has slope $-2$. **c, d** Loss as a function of the fitting parameters – scale factor $\alpha$ and mutation rate $\mu$ – when fitting either the (c) complementary cumulative distribution, or the (d) Laplace transform of the distribution. Plot shows global loss: i.e., summed over $k$. **e, f** Best-fit (e) $\alpha$ and (f) $\mu$ for different values of $k$.

at 30 logarithmically spaced points between $m = 1$ up to the tenth-largest observed mutant count. The loss was defined as the summed squared difference of the $\log_{10}$ CCDF values, pooled across all LPT bin numbers $k$; the best-fit $(\mu, \alpha)$ minimised this loss. As an internal consistency check, the CCDF was also fitted separately for each bin number $k$, see Fig. S4c-f.

As an independent estimator, exploiting a different feature of the data, we fitted the distribution in Laplace space. We evaluated the empirical Laplace transform of the mutant-number distribution,

$$\tilde{P}_{\text{data}}(s) = \frac{1}{\sum_i 1} \sum_i e^{-s\, m_i},$$

at 30 logarithmically spaced values of the transform variable $s$ from $10^{-3}$ to $10^{-1}$, and fitted it to the closed-form theoretical expression derived above and quoted in the main text, $\ln \tilde{P}(s) = \mu N \left(-1 + \mathrm{E}_2(s)\right)$ (Eq. 8 with $\sigma = 1$, $\mathrm{E}_2$ the generalised exponential integral), minimising the summed squared difference in log space over a grid of $\mu$ and $\alpha$.

For the msh2Δ experiment of Fig. 2 of the main text, the global CCDF fit gave a mutation rate $\mu = 2.0 \times 10^{-6}$ per cell per generation with $\alpha = 4.3$; the two estimators and the different bin numbers agreed to within roughly two-fold, with the per-$k$ CCDF fits spanning $1.4–2.9 \times 10^{-6}$ and the Laplace fit giving $2.1–2.3 \times 10^{-6}$. For the three-strain experiment of Fig. 3 of the main text, the fitted mutation rates ordered as expected for the mutator phenotypes, increasing from wild type through msh6Δ to msh2Δ: the global CCDF fits gave $\mu = 2.9 \times 10^{-7}$ for wild-type BY4741, $\mu = 1.0 \times 10^{-6}$ for msh6Δ and $\mu = 3 \times 10^{-6}$ for msh2Δ, with the Laplace-space fit giving consistent per-strain rates ($4.3 \times 10^{-7}$, $1.1 \times 10^{-6}$ and $2.0 \times 10^{-6}$ respectively). Fitted rates are plotted in Fig. S4f and S5e.

In general, we find fitting the Laplace transform more stable to choice of $k$ (see Figs. S4 and S5) and with tighter 95% confidence intervals (see below); in the main text, we quote these fit results for the data in Fig. 3.

### 5.4 Confidence intervals by bootstrapping

We obtained 95% confidence intervals by bootstrapping over barcodes. For each of $B = 10{,}000$ replicates we drew a sample of barcodes with replacement, of the same size as the original set, and repeated the entire analysis on the resampled barcodes. Intervals quoted are plain percentile intervals – the 2.5th and 97.5th percentiles of the replicate distribution. Confidence intervals are in Table 1.

| | $\mu$ ($\times 10^{-6}$) | | $\mu$ per base ($\times 10^{-9}$) | |
|---|---|---|---|---|
| | least squares | Laplace | least squares | Laplace |
| Fig. 2 BY4741 *msh2*Δ | 2.0 [1.5, 2.9] | 2.1 [1.7, 2.4] | 8.6 [6.4, 12.3] | 9.0 [7.1, 10.0] |
| Fig. 3, BY4741 | 0.29 [0.25, 0.37] | 0.43 [0.35, 0.52] | 1.2 [1.1, 1.6] | 1.8 [1.5, 2.2] |
| Fig. 3, BY4741 *msh6*Δ | 1.0 [0.9, 1.4] | 1.1 [0.9, 1.4] | 4.1 [3.6, 6.1] | 4.9 [3.8, 5.9] |
| Fig. 3, BY4741 *msh2*Δ | 3 [2, 10] | 2.0 [1.8, 2.3] | 12 [8, 42] | 8.7 [7.7, 9.8] |

Table 1: Global (all-$k$) mutation-rate fits with 95% percentile bootstrap confidence intervals in brackets, from $10^4$ barcode-level resamples. Per-base values are the same quantities divided by 236, the effective canavanine loss-of-function target size in basepairs [1]. Interval half-widths are rounded to one significant figure and central values to the same decimal place. Each column rounded independently from the unrounded values.

#### 5.5 Data collapse of mismatch repair strains

The canonical Luria-Delbrück distribution depends on the mutation rate and the population size only through their product $\mu N$ – the expected number of independent mutational origins per culture – as is evident from the cumulant generating function (Eq. 8), in which $\mu$ and $N$ enter solely as the scale factor $\mu N$. The three strains that vary in their mismatch repair genotype differ substantially in $\mu$, and so provide a direct test of this prediction: if the mutant-number distribution truly depends only on $\mu N$, then distributions matched in $\mu N$ across strains should collapse onto a single curve.

To match $\mu N$ across strains we used the LPT binning to set each strain's population size. For a chosen target product $P = \mu N$, a strain with best-fit rate $\mu_{\mathrm{strain}}$ requires a bin number $k = \mu_{\mathrm{strain}} N_{\mathrm{total}}/P$; because the strains have different $\mu$, matching a common $P$ requires a different number of bins for each strain. Using the per-strain best-fit $\mu$ obtained above, each strain was binned at its $P$-specific $k$ and its mutant-number CCDF was computed as before, with the absolute scale set by the fitted calibration factor $\alpha$ for each strain. We used the best-fitting values from the fit of the Laplace transform for both $\mu$ and $\alpha$ in this analysis.

We overlaid the three strains' CCDFs for several matched products ($\mu N = 1, 5, 25$) in Fig. 3 of the main text. Within each $\mu N$ family the three strains' distributions collapse onto the common theoretical curve, while increasing $\mu N$ shifts the whole family up and to the right — confirming that, once the mutation rate has been accounted for, the mutant-number statistics of the wild-type and mutator strains are described by the same canonical Luria-Delbrück distribution.

#### 5.6 Maximum likelihood inference of induced mutagenesis in UV-exposed populations

Two of the three populations analysed in Fig. 4 of the main text were exposed to UV immediately prior to selection (*UV exposure*, above). Mutations induced by that exposure should differ from spontaneous ones in a way that is visible in the mutant-number distribution: because they arise at the very end of the growth period, the cells carrying them have no time to divide again, and so each induced mutation contributes exactly one mutant rather than a clone of $e^{\lambda(T-t)}$ mutants. Induced mutagenesis therefore adds no jackpots, and instead contributes a simple Poisson component. Writing $\mu$ for the spontaneous rate and $\mu_{\mathrm{ind}}$ for the induced rate per cell, the number of mutants in a lineage of $N$ cells is

$$m = m_{\mathrm{LD}} + m_{\mathrm{ind}}, \qquad m_{\mathrm{ind}} \sim \mathrm{Poisson}\left(\mu_{\mathrm{ind}} N\right), \tag{11}$$

where $m_{\mathrm{LD}}$ is drawn from the Luria-Delbrück distribution for rate $\mu$ and population size $N$. Setting $\mu_{\mathrm{ind}} = 0$ recovers the purely spontaneous model, so the two models are nested and can be compared directly.

We first fixed the scale-factor $\alpha$ relating reads to absolute number of mutants by fitting the unexposed (i.e. no-UV) population to the canonical (spontaneous-mutations only) Luria-Delbrück distribution by the least-squares CCDF fit described above; the per-$k$ estimates varied between 3.0 and 4.8, and we took their median, $\alpha = 3.4$, as the value for all three populations in this experiment.

Fitting data by maximum likelihood requires the probability distribution of $m$, which has no closed form and was therefore built numerically. For each population, bin number $k$ and spontaneous rate $\mu$, we simulated $3 \times 10^6$ replicates of the Luria-Delbrück process by the algorithm of *Numerical calculations of the Luria-Delbrück distribution*. Re-using that same set of replicates, we then added an independent Poisson($\mu_{\mathrm{ind}} N$) draw for each value of $\mu_{\mathrm{ind}}$ on the grid, and histogrammed the resulting mutant numbers onto a fixed, fine set of bin edges – one bin covering $[0, 0.5)$ together with 300 logarithmically spaced edges from 0.5 to $10^{12}$ – to obtain a numerical probability distribution. Interpolating the cumulative sum of this distribution gives

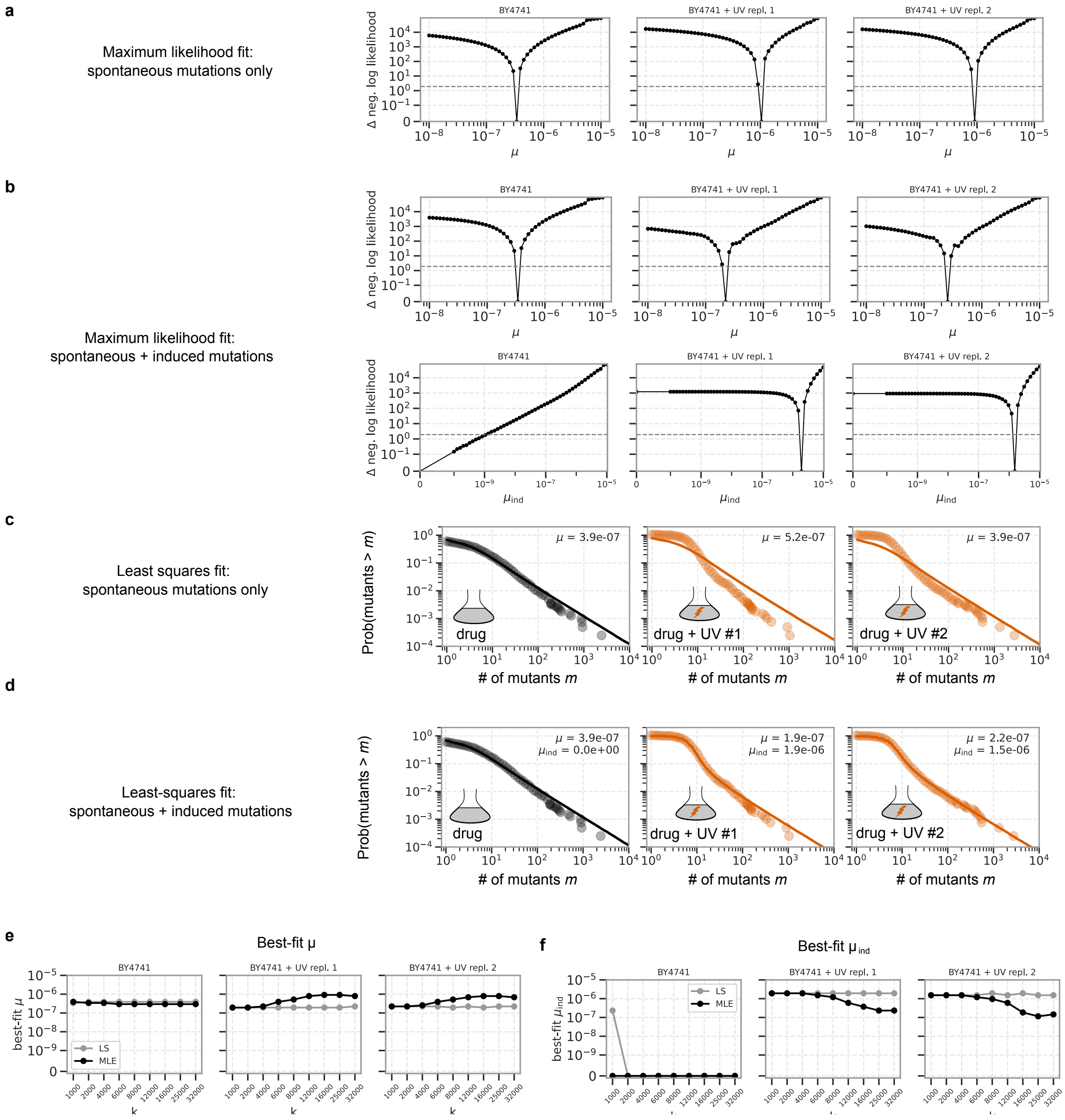

Figure S6: **Maximum likelihood fitting of data in Fig. 4 of the main text.** **a** The deviation in log-likelihood from the best-fit $\mu$, as a function of $\mu$, for all three datasets when they are fit to a model with only spontaneous mutagenesis. **b** As in (a), but for $\mu$ and $\mu_{\text{ind}}$ from a fit to a model with both spontaneous and induced mutagenesis. **c, d** Least-squares fit of the CCDF to a model with only (c) spontaneous or (d) spontaneous + induced mutagenesis. **e, f** Best-fit (e) $\mu$ and (f) $\mu_{\text{ind}}$ for different values of $k$, for both least-squares (LS) and maximum likelihood (MLE) fitting. All results in (a) through (d), and in Fig. 4 of the main text, are shown for $k = 4000$.

both the CCDF at any mutant number and the probability of any coarser bin, so that a single simulation per $(\mu, \mu_{\text{ind}}, k)$ serves both the likelihood fit described here and the least-squares fit described below. Since $\alpha$ rescales only the data, the simulations are independent of it and were computed once.

To evaluate the likelihood, the $k$ observed mutant numbers were discretised onto coarse bins: one bin covering $[0, 1)$, then 50 logarithmically spaced bins up to twice the largest observed mutant number. Writing $b(i)$ for the bin containing lineage $i$ and $P_b$ for the model probability of bin $b$, the log-likelihood is

$$\ln \mathcal{L}\left(\mu, \mu_{\text{ind}}\right) = \sum_{i=1}^{k} \ln P_{b(i)}\left(\mu, \mu_{\text{ind}}\right), \tag{12}$$

which we maximised by exhaustive evaluation on a grid of 50 logarithmically spaced values of $\mu$ between $10^{-8}$ and $10^{-5}$, and of $\mu_{\text{ind}} = 0$ together with 50 logarithmically spaced values between $10^{-10}$ and $10^{-5}$. $P_{b(i)}\left(\mu, \mu_{\text{ind}}\right)$ was obtained from the simulations described above.

Fitting the spontaneous-only model, the negative log-likelihood rises steeply within a few grid steps of its optimum for all three populations (Fig. S6a), as it does for the combined model when each parameter is profiled over the other (Fig. S6b). For the unexposed population the likelihood was maximised at $\mu_{\text{ind}} = 0$, with a spontaneous rate of $\mu = 3.4 \times 10^{-7}$. Both exposed populations instead required a substantial induced component: $\mu_{\text{ind}} = 1.9 \times 10^{-6}$ with $\mu = 2.2 \times 10^{-7}$ for UV population 1, and $\mu_{\text{ind}} = 1.5 \times 10^{-6}$ with $\mu = 2.6 \times 10^{-7}$ for UV population 2. The spontaneous rates recovered for the exposed populations are of the same order as that of the unexposed population. Forcing $\mu_{\text{ind}} = 0$ for the exposed populations instead inflates the apparent spontaneous rate to approximately $10^{-6}$ (Fig. S6a).

As a check that these conclusions do not depend on the likelihood construction, we repeated the fit by least squares, minimising the same log–log CCDF loss used above over $\mu$ only or $(\mu, \mu_{\text{ind}})$ with $\alpha$ fixed. Again, the spontaneous-only model fails to fit the UV data well – though with a different inferred $\mu$ compared to maximum likelihood, Fig. S6c, reflecting the different weighting of features in the data by different fitting methods. When fitting the spontaneous + induced model, the obtained mutation rates are similar to those obtained by maximum likelihood, and the least-squares fit of the combined model reproduces the observed distributions across their full range (Fig. S6d).

Finally, the fits were repeated for bin numbers from $k = 1000$ to $k = 32{,}000$ (Figs. S6e, f). We find that the least-squares fit is broadly insensitive to the choice of $k$, while the maximum likelihood estimates begin to diverge as $k$ becomes large. A potential explanation is that, as $k$ increases, the population size $N$ per virtual lineage decreases ($N \sim 10^{10}/k$, where $10^{10}$ is the total number of cells in the flask pre-selection). The number of barcodes that do not get any mutants is $P(m = 0) = \exp\left(-\mu N - \mu_{\text{ind}} N\right) \approx e^{-10^{-6} N}$ – which becomes appreciable as $k$ crosses $10^4$. At large $k$, then, the bulk of the likelihood is decided by lineages that produce no mutants – which cannot, by nature, distinguish between spontaneous and induced mutations. In contrast, the least-squares fit is sensitive to the tail of the distribution – which continues to be informative at larger $k$. All values quoted above, and all panels of Fig. 4 of the main text, use $k = 4000$, where both fits agree.

To obtain an independent estimate of $\mu_{\text{ind}}$, we plated 1mL of the t=0 cultures on canavanine before and after exposure to UV. For UV population 1, we counted 529 colony forming units (cfus) prior to UV, and 726 after. For UV population 2, we counted 83 cfus before, and 219 cfus after. From this, we estimated that UV exposure lead to $2.6 \times 10^4$ and $1.8 \times 10^4$ new mutants in UV populations 1 and 2, respectively. Assuming a total population size of $1.2 \times 10^{10}$ prior to selection,

$$\mu_{\text{ind}} = \begin{cases} \frac{2.6\times10^4}{1.2\times10^{10}} \approx 2.2 \times 10^{-6} & \text{for UV population 1} \\ \frac{1.8\times10^4}{1.2\times10^{10}} \approx 1.5 \times 10^{-6} & \text{for UV population 2} \end{cases}$$

which is comparable to the value of $\mu_{\text{ind}}$ inferred by maximum likelihood estimation.

#### 5.7 Reanalysing Fig. 2 data to simulate a ‘D$\mu$S’

In Fig. 5 of the main text, we used the data from Fig. 2 to determine the feasibility of a high-throughput ‘deep mutation rate scan’ (D$\mu$S). To do so, we partitioned the 119,672 barcodes from that experiment into sets of 50, 100, 200, or 400 barcodes. For each set, the mutation rate was extracted by a simple maximum likelihood generalisation of the traditional $P_0$ method [3]. In brief, we first compute the indicator variable $s_i$ for every barcode, where $s_i = 1$ if $m_i >= 1$ and 0 otherwise, where $m_i$ is the number of mutants produced by the barcode. To obtain $m_i$ from read counts, we used the calibration factor $\alpha$ determined above (see Fig. S4 and related text). In terms of $s_i$, we define the likelihood function:

$$\mathcal{L} = \prod_{\text{bcds } i \in \text{ set}} \left(e^{-\mu N_i}(1 - s_i) + (1 - e^{-\mu N_i})s_i\right)$$

where $N_i$ is the pre-selection population size. The likelihood was maximised over $\mu$ for each set, and the histogram in Fig. 5 of the main text constructed over sets.
